# Efficient *in situ* epitope tagging of rice genes by nuclease-mediated prime editing

**DOI:** 10.1101/2024.07.29.605657

**Authors:** Xueqi Li, Sujie Zhang, Chenyang Wang, Bin Ren, Fang Yan, Shaofang Li, Carl Spetz, Jinguang Huang, Xueping Zhou, Huanbin Zhou

## Abstract

*In situ* epitope tagging is crucial for probing gene expression, protein localization, and the dynamics of protein interactions within their natural cellular context. However, the practical application of this technique in plants presents considerable hurdles. Here, we comprehensively explore the potential of the CRISPR/Cas nuclease-mediated prime editing and different DNA repair pathways in epitope tagging of endogenous rice genes. We find that SpCas9 nuclease/microhomology-mediated end joining (MMEJ)-based prime editing strategy (termed NM-PE) facilitates more straightforward and efficient gene tagging compared to the conventional and other derivative PE method. Furthermore, the PAM-flexible SpRY and ScCas9 nucleases-based prime editors have been engineered and implemented for the tagging of endogenous genes with diverse epitopes, significantly broadening the applicability of NM-PE in rice. Moreover, NM-PE has been successfully adopted in simultaneous tagging of *OsMPK1* and *OsMPK13* in rice plants with c-Myc and HA tags, respectively. Taken together, our results indicate great potential of the NM-PE toolkit in the targeted gene tagging for Rice Protein Tagging Project, gene function study and genetic improvement.

## Introduction

The utilization of transgenes coupled with epitope tags (e.g., FLAG, HA, c-Myc, *etc*.) have been widely employed in elucidating the underlying biological mechanisms associated with agronomic traits such as yield, quality, and resistance to biotic and abiotic stresses (Yusibov et al. 2016). However, the random integration of transgenes often fails to fully recapitulate the dynamic expression profiles of endogenous genes, and this limitation is not addressed by driving transgene expression with native promoters, as they may involve long-range chromatin interactions between proximal and distal regulatory elements (Li et al. 2019). Also, overexpression of a gene can sometimes lead to artifacts, which do not reflect the natural function of the gene in its normal context (Gibson et al. 2013). Over the past decade, the advent of genome editing technologies has revolutionized the field of functional genomics and has brought significant improvements to *in situ* epitope tagging techniques, which allows for the expression of a tagged gene under the control of its endogenous promoter and regulatory elements (Lu et al. 2020a).

Genome editing tools, especially the Clustered Regularly Interspaced Short Palindromic Repeats (CRISPR)/CRISPR-associated nucleases 9 (Cas9) system, have been utilized for site-specific genetic modifications in a wide range of plant species, including rice, wheat, maize, soybean, *etc* (Zhou et al. 2014; Zhu et al. 2020; Gao 2021). The simplified system consists of the endonuclease protein Cas9 and a single guide RNA (sgRNA), enabling it to cleave genomic DNA at a specific site and result in DNA double-strand breaks (DSBs) (Liu et al. 2022). The DSBs can be repaired with or without exogenous DNA donor templates through different cellular DNA repair pathways, such as non-homologous end joining (NHEJ), microhomology-mediated end joining (MMEJ), homology-directed repair (HDR), and resulting in targeted DNA fragment insertion and replacement (Nambiar et al. 2022). As a proof of concept, CRISPR/Cas9 nuclease-mediated targeted epitope tag insertions were successfully achieved in rice plant by utilizing NHEJ-TR-HDR approach with chemically modified blunt-ended double-stranded oligodeoxynucleotides (dsODNs) (Lu et al. 2020b). This approach was further simplified by 1-nt complementary 5’ overhang dsODNs, which harnesses the staggered cleavage activity of CRISPR/Cas9 and facilitates directional targeted insertion of epitope tag in green foxtail protoplasts by NHEJ only (Kumar et al. 2023). Nevertheless, CRISPR/Cas nuclease-based strategies for plant gene tagging necessitate the use of exogenous DNA donor, leading to issues of inconvenience, high costs, and low efficiency.

In recent years, significant progress has been made in the development of prime editing (PE), which enables targeted insertions, deletions, and all 12 types of point mutations without exogenous donor DNA repair templates (Anzalone et al. 2019). Prime editors consist of the nickase Cas9 (H840A), a Moloney-murine leukemia virus reverse transcriptase (M-MLV RT), and a PE guide RNA (pegRNA) (Anzalone et al. 2019). The pegRNA contains an RT template (RTT) encoding intended edits and a primer binding site (PBS) for hybridization of the 3’ end of the nicked DNA strand to initiate reverse transcription. After the reverse transcription, the desired sequence is ultimately integrated into the target site through a series of complex DNA repair mechanisms, including 3’ flap annealing, 5’ flap displacement, 5’ flap excision, and heteroduplex resolution (Chen and Liu 2023). The PE has been initially characterized by limited editing efficiency in plants (Hua et al. 2020; Lin et al. 2020; Tang et al. 2020). To address this issue, numerous endeavors have been directed towards optimizing PE, including employing engineered PE proteins to enhance their binding and reverse transcription activities (Chen et al. 2021; Li et al. 2022b; Zong et al. 2022; Doman et al. 2023; Ni et al. 2023); using a composite promoter to increase pegRNA transcription (Jiang et al. 2020; Jiang et al. 2022b); optimizing pegRNA reverse transcription efficiency by modifying the PBS melting temperature (Lin et al. 2021); incorporating a 3’ structural RNA motif to reduce pegRNA degradation (Nelson et al. 2022); incorporating additional silent mutations near the intended edit to avoid recognition by the DNA mismatch repair (MMR) pathway (Chen et al. 2021; Xu et al. 2022); employing paired pegRNAs to facilitate the precise insertion of edits (Lin et al. 2021; Anzalone et al. 2022; Wang et al. 2022a; Liu et al. 2024); introducing an additional nick in the unedited strand (Xu et al. 2022); and using a surrogate system to enrich the editing events (Li et al. 2022a). The continuous optimization of PE is positioned to significantly advance the development of *in situ* epitope tagging in plants. For example, the original version of prime editor (PE3) was successfully employed to introduce the HA epitope tag to the endogenous *CCA1* gene in *Arabidopsis* as well as FLAG tag to *OsSULTR3;6* in rice, albeit with notably low efficacy (Wang et al. 2021b; Hong et al. 2024). Equipped with a viral nucleocapsid (NC) protein, a 3’ structural RNA motif, *etc*., the updated PE system (ePE2) achieved effective tagging, ranging from 18.75-50.00%, at the *OsUBQ10* and *OsTubA1* loci in transgenic rice plants (Li et al. 2023a). Moreover, application of dual pegRNAs with ePE2 has been shown to enhance PE efficiency, enabling 66-nt 3xFLAG tag replacement. Lately, NEPE, characterized by M-MLV RT at the N-terminus of Cas9 (H840A), has been reported to achieve a tagging efficiency of 20.00% for the FLAG tag and 15.00% efficiency for the 6xHis tag when targeting *OsMSP* in rice plants (Zhong et al. 2024). Anyhow, PE through *Agrobacterium*-mediated transformation offers a cost-effective and practical approach for endogenous gene tagging in plants. Nevertheless, the constrained efficacy of PE-mediated gene tagging somewhat limits its application.

To date, various derivative PE strategies with diverse molecular mechanisms have been also explored to achieve precise sequence insertion and replacement, such as Cas12a-mediated REDRAW editing and CPE (Bill Kim et al. 2022; Liang et al. 2024), DNA-dependent DNA polymerase-mediated Click editing (Zheng et al. 2023; Ferreira da Silva et al. 2024), template-jumping PE (Liu et al. 2023), *etc*. Considering the great power of CRISPR/Cas nuclease to induce DSBs at the target sites in plant cells, it can activate highly efficient NHEJ and/or MMEJ pathways, facilitating targeted insertion or deletion of genomic fragments (Tan et al. 2020; Wang et al. 2024). Therefore, the nuclease-mediated derivative PE might provide additional gene tagging strategies, potentially with improved editing efficiency. Here, we have assessed the feasibility of utilizing NHEJ and MMEJ pathways in CRISPR/Cas nuclease-mediated PE for epitope tagging of endogenous genes in rice. Our findings reveal that both DNA repair pathways can be employed to achieve *in situ* epitope tagging, with the SpCas9 nuclease/MMEJ-based prime editing strategy (NM-PE) outperforming the conventional PE method. Furthermore, NM-PE has been effectively utilized in single and dual gene tagging with various epitope tags in rice plants. Moreover, our study demonstrates that the PAM-flexible SpRY and ScCas9 nucleases are suitable for NM-PE as well, which significantly expand the scope of gene tagging in the rice genome. All the data implicate the great potential of applying the NM-PE toolkit in endogenous gene tagging in plants.

## Results

### The establishment of a high-efficiency prime editing system in rice

To develop a high-efficiency PE system in rice, we have adopted most strategies reported for the optimization of PE to date (Lin et al. 2021; Jiang et al. 2022b; Li et al. 2022b; Nelson et al. 2022; Xu et al. 2022; Zong et al. 2022). The coding sequences of the SpCas9n (R221K/N394K/H840A) variant and the Moloney murine leukemia virus reverse transcriptase pentamutant without the RNase H domain (M-MLV RT-ΔRNase H/D200N/L603W/T330P/T306K/W313F) were codon-optimized for expression in rice. The M-MLV RT mutant was fused to the C-terminus of SpCas9n variant with multiple copies of bipartitle nucleus localization signal (bpNLS) and c-Myc NLS incorporated at both terminus and in the linker, respectively (Supplementary Figure S1A). The chimeric gene, named *rPE14a*, was used to replace the *SpRY* gene in the binary vector pUbi:SpRY (Xu et al. 2021), resulting in pUbi:rPE14a in which *rPE14a* is under the control of the maize ubiquitin 1 promoter. Meanwhile, an engineered pegRNA module that includes ribozyme and tRNA processing systems, and a structured RNA motif of modified prequeosine_1_-1 riboswitch aptamer (evopreQ1) was designed and inserted downstream of the CaMV 35S enhancer-CmYLCV-U6 composite promoter in pENTR:sgRNA41. Thus, the pegRNA-expressing cassette can be transferred into the pUbi:rPE14a via Gateway recombination and used for *Agrobacterium*-mediated rice transformation (Wang et al. 2022b).

Next, we designed a pegRNA composed of a 10-nt PBS (30℃ melting temperature) and a 11-nt of RTT for changing TG<u>G</u> (W) to TG<u>C</u> (C) at OsACC-W2125 site in the rice genome and used it to evaluate the editing capability of rPE14a in transgenic rice plants (Supplementary Figure S1B). After transformation, PCR amplicon of the target region from T0 independent transgenic lines was subjected to next-generation sequencing (NGS) and further randomly verified by Sanger sequencing. We found that 95.83% (69 out of 72 lines) of hygromycin-resistant lines carried the desired G-to-C transversion at *OsACC* site. Among these, 64 were homozygous mutants, two were heterozygous, and three had biallelic mutations that carried the desired edit and unintended byproducts (Supplementary Figure S1C-E). These data indicate that rPE14a we engineered is a high-activity rice prime editor.

### PE-mediated *in situ* epitope tagging in rice

Considering the high editing efficiency of rPE14a, we investigated its potential for application in precisely epitope tagging of endogenous rice genes (Figure 1A, B). A pegRNA was designed to incorporate the FLAG tag immediately upstream of the *OsMPK1* stop codon. It featured an 11-nt PBS with a melting temperature of 30℃ and a 44-nt RTT. The RTT included two additional nucleotides which kept FLAG tag in frame with *OsMPK1*, and a 15-nt homology arm with two synonymous mismatches (Figure 1C). Another pegRNA was designed for *OsMPK13* using the same strategy, which contained a 10-nt PBS and a 44-nt RTT (Figure 1D). In T0 plants, approximately 21.74% (10 out of 46 lines) of independent transgenic lines exhibited the desired *in situ* FLAG tagging of *OsMPK1*, all of which carried heterozygous tag insertions (Figure 1E). As of *OsMPK13*, precisely FLAG tagging occurred in 23.91% (11 out of 46 lines) of lines, including seven with heterozygous insertions and four with biallelic edits (Figure 1E). Detailed analysis of the edited alleles revealed that approximately 11% of the DNA double-strands at both target sites experienced precise insertions of FLAG tag sequences (Figure 1F). To be mentioned, 14 and 26 independent lines harbored byproducts in *OsMPK1* and *OsMPK13*, respectively. In addition to Indels, most detected byproducts were imprecise edits, involving either incomplete insertion epitope tags or additional flanking sequences (Supplementary Figure S2A, B) Collectively, these data suggest that the high-efficiency rPE14a-mediated prime editing enables *in situ* epitope tagging of rice genes through targeted insertion with reasonable efficiency.

**Figure 1.**
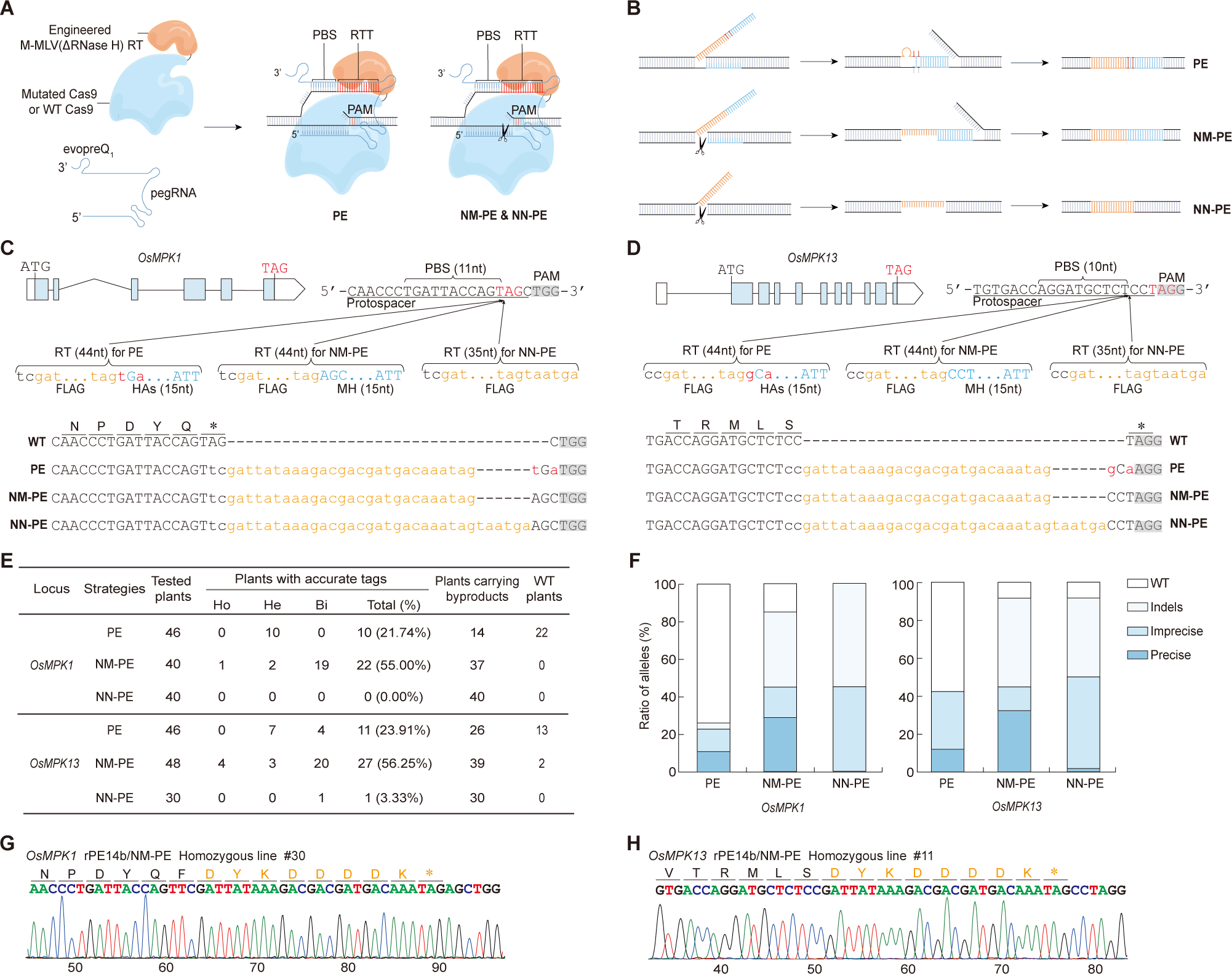
*In situ* epitope tagging in rice using different PE strategies. **A**) A schematic illustration depicts various prime editors for endogenous gene tagging. The epitope tag sequence, embedded within the reverse transcriptase templates (RTT) of evopreQ1-modified pegRNA, is inserted at the target site in the rice genome. PBS, primer binding site; PAM, protospacer adjacent motif; PE, conventional prime editing; NM-PE, nucleases/MMEJ-based prime editing; NN-PE, nucleases/NHEJ-based prime editing. **B**) Schematics outline the gene tagging mechanisms facilitated by different PE strategies. Following reverse transcription, the epitope tag sequence is integrated into the target site through various DNA repair pathways. The sequences of insertion, homology, and mismatches are highlighted in orange, blue, and red, respectively. **C**) Three pegRNAs were designed to incorporate the FLAG epitope immediately upstream of the *OsMPK1* stop codon with different PE strategies. Exons are represented as blue boxes, and introns as black lines. The PAM sequences and insertion sites are highlighted in gray and indicated with arrows, respectively. Inserted sequences are displayed in lowercase. Homology arm (HAs) with two synonymous mismatches and microhomology arm (MH) are noted. **D**) As for (**C**), but showing *in situ* epitope tagging at the *OsMPK13* site. **E**) Summary of the frequencies of gene-tagging created by PE, NM-PE, and NN-PE in T0 transgenic rice lines. Plants with accurate tags exhibit at least one allele at the target site with an exact insertion, categorized as homozygous (Ho), heterozygous (He), and biallelic (Bi) where precise edits and byproducts reside on different chromosome. Conversely, plants with byproducts display at least one allele at the target site with imprecise insertions and/or Indels. **F**) The stacked diagram illustrates the proportions of distinct editing outcomes in alleles created by PE, NM-PE, and NN-PE. **G-H**) Sanger sequencing chromatograms demonstrate in-frame FLAG insertions at the *OsMPK1* (**G**) and *OsMPK13* (**H**) sites in representative T0 lines, respectively. In (**C-D**) and (**G-H**), amino acid sequences are annotated above corresponding DNA sequences.

### Nuclease-mediated PE for *in situ* epitope tagging in rice

To explore enhancing the efficiency of PE-mediated endogenous gene tagging, we sought to develop derivative PE systems that utilize nuclease-mediated PE and different DSB repair mechanisms in rice. We designed two distinct strategies: the **n**uclease/**M**MEJ-mediated **PE** (NM-PE) and the **n**uclease/**N**HEJ-mediated **PE** (NN-PE) (Figure 1A, B). We hypothesized that the SpCas9/pegRNA complex would precisely cleaves the target site, enabling the loading of the epitope tag sequence at the DSB by M-MLV RT through either *in vivo* MMEJ or NHEJ pathways in the presence and absence of a microhomology (MH) arm, thereby achieving epitope tagging of target genes in the rice genome. Accordingly, we reverted the SpCas9 (R221K/N394K/H840A) in rPE14a to its wild-type version, resulting in rPE14b. Both *OsMPK1* and *OsMPK13* were then targeted again for FLAG tag insertion in transgenic plants using rPE14b, with or without 15-nt MH arms (Figure 1C, D).

After transformation, the NM-PE strategy accomplished precisely epitope tagging in 22 out of 40 independent lines (55.00% efficiency) at the *OsMPK1* site, a 2.53-fold increase compared to rPE14a-mediated PE. At the *OsMPK13* site, NM-PE achieved an efficiency of 56.25% (27 out of 48 lines), a 2.35-fold increase over PE. Notably, 2.50% and 8.33% of plants displayed homologous insertions at the *OsMPK1* and *OsMPK13* sites, respectively (Figure 1E, G, H). Further analysis showed that 28.75% of *OsMPK1* alleles and 32.29% of *OsMPK13* alleles carried intact FALG tag, representing 2.64- and 2.70-fold improvement over PE at both tested sites, respectively (Figure 1F). These data indicate that the NM-PE strategy significantly enhances *in situ* epitope tagging in rice. Beyond the imprecise tagging, Indel mutations, which resulted from the anticipated high nuclease activity of rPE14b, were frequently observed in byproducts (Figure 1F; Supplementary Figure S2C, D). Regarding NN-PE, only 1 line with biallelic edits was identified with precise sequence insertion at the *OsMPK13* site (Figure 1E). Instead, 45.00% of *OsMPK1* and 48.33% of *OsMPK13* alleles carried imprecise sequence insertions (Figure 1F). To be mentioned here, three consecutive stop codons (<u>TAG</u>TAA<u>TGA</u>) were included in RTTs to ensure the presence of termination signals in target genes, even in cases of nucleotide deletion occurred in NN-PE (Supplementary Figure S2E, F; Figure S3). As a result, 22.50% and 43.33% of lines were considered as wanted plants with epitope tagging of *OsMPK1* and *OsMPK13*, respectively (Supplementary Figure S3C). Thus, NN-PE is capable of endogenous gene tagging in rice as well. Combined, these results suggest that NM-PE outperforms the conventional PE and NN-PE in *in-situ* epitope tagging in the rice genome.

Next, we further verified the feasibility of NM-PE by tagging the endogenous *OsMPK8* gene (Supplementary Figure S4A). Out of 48 independent lines, 19 lines (39.58% efficiency) exhibited the precise insertion of the FLAG tag at the target site (Supplementary Figure S4B), of which 2 were homologous and 19 carried biallelic edits (Supplementary Figure S4C). Additionally, 44 lines were identified with byproducts, harboring either Indels or truncated FLAG tag insertions (Supplementary Figure S4D). Collectively, we conclude that NM-PE is a straightforward and efficient strategy for *in situ* epitope tagging of rice genes.

### Broadening the targeting scope of NM-PE using PAM-flexible Cas9 variants

Given the stringent NGG protospacer adjacent motif (PAM) requirement for SpCas9-based rPE14b in targeted gene tagging, we explored the use of PAM-flexible Cas nucleases to enhance NM-PE (Chatterjee et al. 2018; Ren et al. 2019; Walton et al. 2020; Wang et al. 2022c), aiming to expand the range of targetable genes throughout the rice genome. To this end, we upgraded rPE14b by substituting SpCas9 with the SpRY nuclease, which is capable of recognizing NRN and certain NYN PAMs in rice (Xu et al. 2021), resulting in rPE15b. Next, we designed three pegRNAs to introduce FLAG, HA, and c-Myc epitope tags to the 3’ ends of *OsMPK3*, *OsMPK4*, and *OsMPK11*, respectively, utilizing TGA and AGA PAMs located at the stop codon sites. To ensure the integrity of the C-terminals of MPKs and prevent the target sites from being re-cut in gene-tagged plants, 2-6 additional nucleotides (corresponding to an additional 0-1 amino acid) beyond the epitope tag sequences were introduced into RTT as well (Figure 2A-C). After genotyping T0 transgenic rice plants, we identified only 1 line (2.38%, biallelic) with FLAG-tagged *OsMPK3* and 3 (6.25%) with c-Myc-tagged *OsMPK11*. Among the *OsMPK11*-tagged lines, 2 were heterozygous and 1 carried biallelic edits (Figure 2D-F). Regarding *OsMPK4*, no HA-tagging plants were isolated. Of note, 6, 15, and 24 lines were identified with byproducts at the *OsMPK3*, *OsMPK4*, and *OsMPK11* sites, respectively. Further analysis indicated that the nuclease activity of SpRY accounted for 8.33% to 40.00% of mutations (Figure 2G-J). All these data imply that the three target sites remain accessible for SpRY-based rPE15b, albeit with low gene tagging efficiency. It has been reported that the broad PAM compatibility of the SpRY nuclease enables self-editing of the sgRNA region within the transferred T-DNA in transgenic rice cells (Xu et al. 2021). Thus, rPE15b-mediated epitope tagging may also be susceptible to self-editing, which could disrupt the pegRNA and thereby reduce the efficiency of tag insertion. Given this consideration, we investigated the integrity of pegRNA transgenes in all T0 transgenic plants. We found that self-editing of pegRNAs within the T-DNA was highly induced, ranging from 33.33% to 64.58% (Figure 2F, K). Collectively, these results indicate that the suboptimal performance of SpRY-mediated NM-PE in endogenous gene tagging may be attributable to both its site-dependent recognition of non-canonical PAMs and its propensity for self-editing.

**Figure 2.**
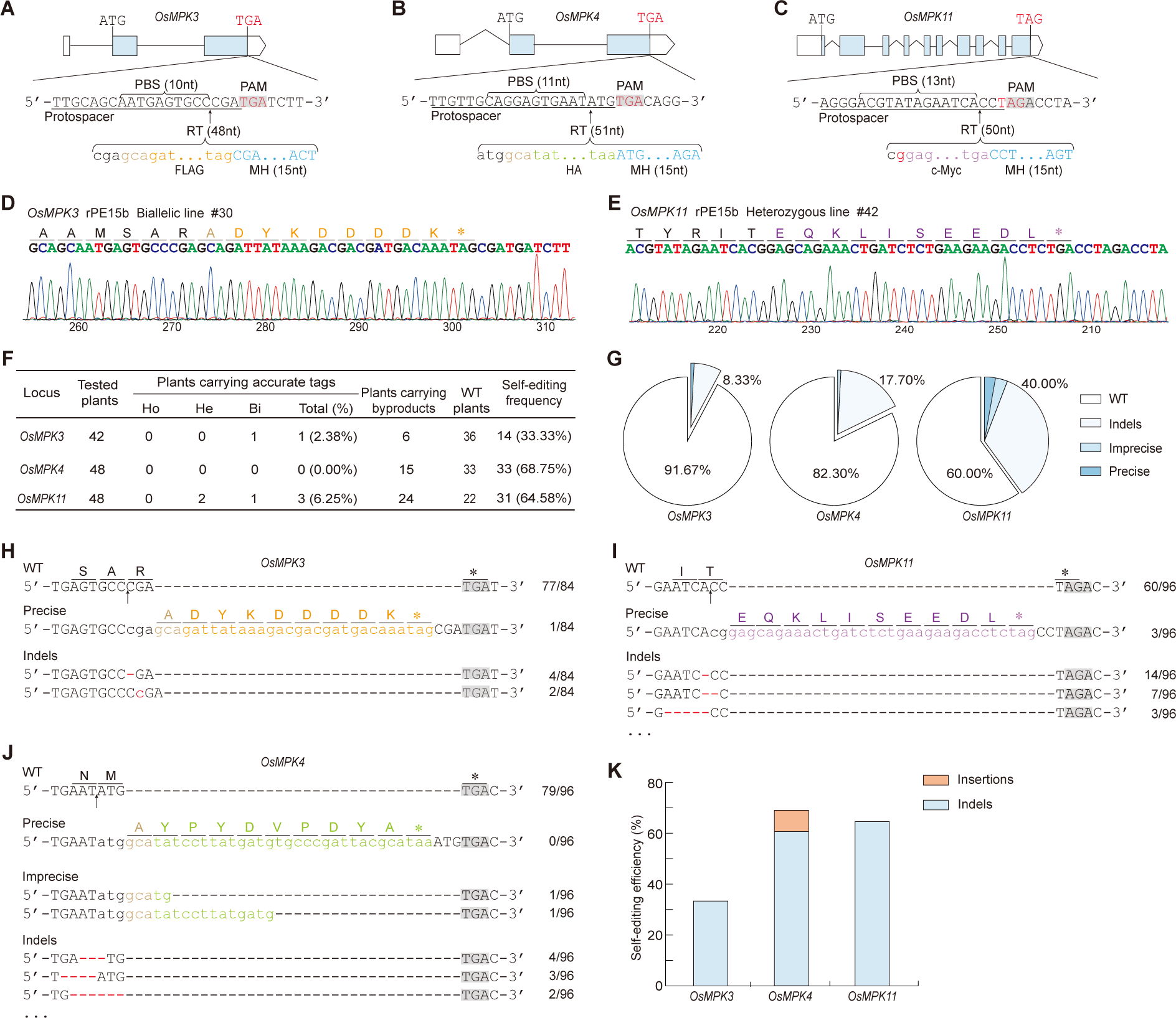
*In situ* epitope tagging through SpRY-mediated NM-PE in rice. **A-C**) The pegRNAs were designed to insert FLAG epitope tag at the *OsMPK3* site (**A**), HA epitope tag at the *OsMPK4* site (**B**), and c-Myc epitope tag at the *OsMPK11* site (**C**) through SpRY-mediated NM-PE (rPE15b). To ensure the C-terminal integrity of the MPKs and prevent target site from being re-cutting by SpRY nuclease, 2-6 additional nucleotides were incorporated into the RTTs beyond the epitope sequences. **D**) Representative Sanger sequencing chromatograms display *in situ* FLAG epitope tagging at the *OsMPK3* site. TA cloning was performed on PCR products from these targeted regions for sequencing. **E**) As for (**D**), but showing c-Myc epitope tag at the *OsMPK11* site. **F**) Summary of the frequencies of gene tagging and self-editing created by rPE15b in T0 transgenic rice lines. Plants with precise tags exhibit at least one allele at the target site with an exact insertion, categorized as homozygous (Ho), heterozygous (He), and biallelic (Bi) where edits and byproducts reside on different chromosome. Conversely, plants with byproducts display at least one allele at the target site with imprecise insertions and/or Indels. **G**) Pie charts depict the distribution of editing types in alleles of T0 mutants. **H-J**) Representative alignment and allele frequencies of the *OsMPK3* site (**H**), the *OsMPK11* site (**I**), and the *OsMPK4* site (**J**) created by rPE15b in T0 transgenic rice lines. **K**) The efficiency of various self-editing events induced by rPE15b. In (**A-C**) and (**H-J**), PAM and insertion sites are highlighted in gray and indicated with arrows, respectively. Deletions are marked by red dashed lines. Inserted sequences are displayed in colored lowercase letters: brown for sequences designed to prevent re-cutting at target sites, orange for the FLAG epitope, purple for the c-Myc epitope, and light green for the HA epitope.

Therefore, we switched to the CRISPR/ScCas9 system (Figure 3A), which recognizes the minimal NNG PAM for genome editing without self-editing activity, and generated derivative prime editor rPE16b (Wang et al. 2020). The endogenous genes *OsMPK6*, *OsMPK7*, and *OsMPK10* were tested by FLAG and c-Myc tagging towards GAG and TAG PAM at the stop codon sites, respectively (Figure 3B-D). We found that ScCas9-based rPE16b resulted in precise FLAG tagging in 20.00% (4 out of 20) of lines for *OsMPK6*, 70.83% (34 out of 48) of lines for *OsMPK10*, and precise c-Myc tagging in 62.22% (28 out of 45) of lines for *OsMPK7* (Figure 3E-H). Notably, homozygous gene tagging occurred in 22.22% of plants at the *OsMPK7* site and 25.00% at the *OsMPK10* site (Figure 3H). We assume that the low efficiency of tagging *OsMPK6* could be attributable to the site-dependent recognition of GAG PAM by ScCas9. Alternatively, high frequencies of biallelic edits and Indel mutations were discovered. Detailed analysis of the edited alleles revealed that the ScCas9-mediated NM-PE editing paradigm largely resembles the performance of spCas9 version mentioned above, albeit discernible site-specific distinctions were noted (Figure 3I; Supplementary Figure S5).

**Figure 3.**
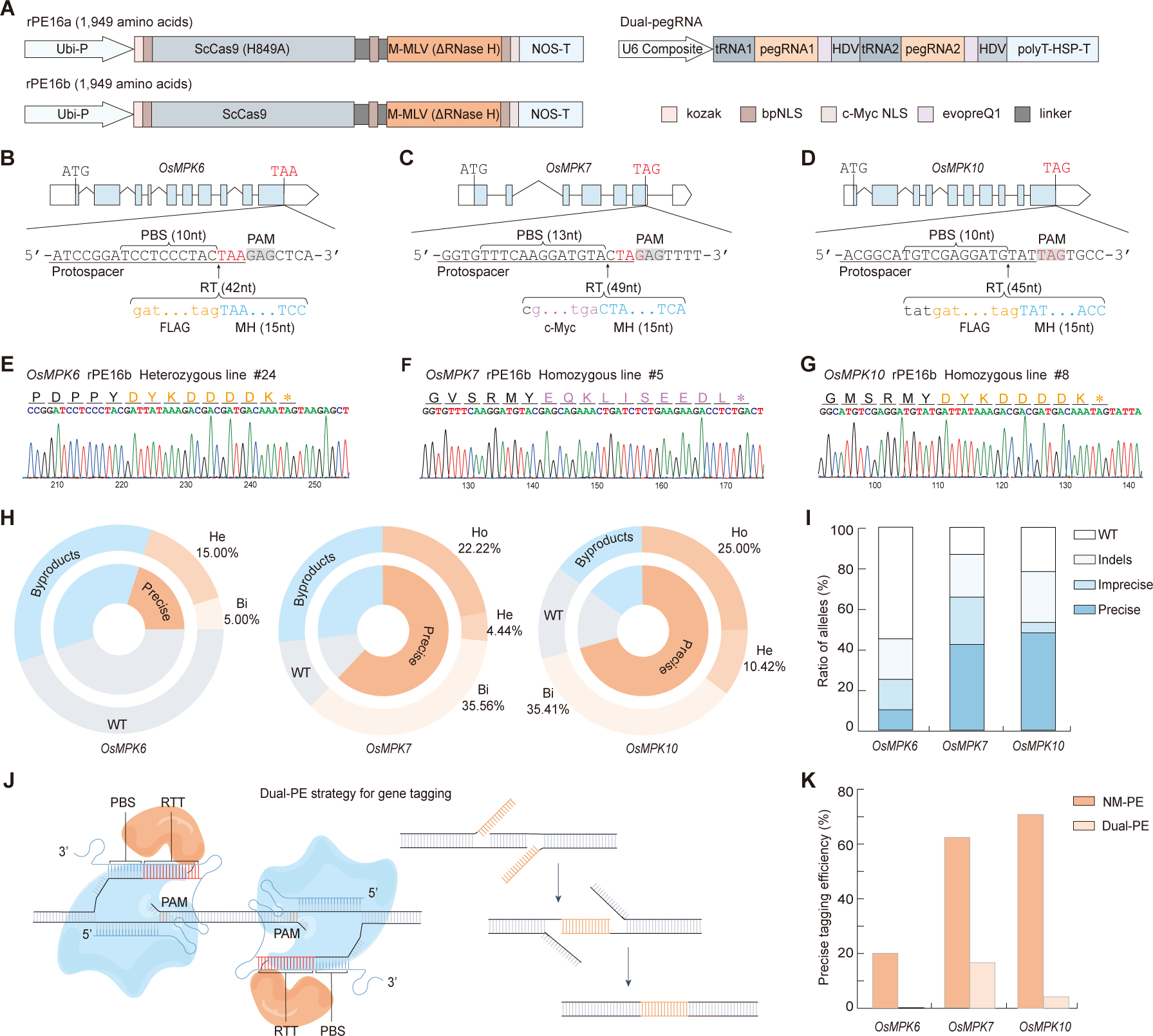
*In situ* epitope tagging through ScCas9-mediated NM-PE in rice. **A**) Schematic of the rPE16a, rPE16b, and dual-pegRNA vectors: rPE16a is designed for gene tagging with dual pegRNA (Dual-PE), while rPE16b is optimized for gene tagging via NM-PE. tRNA1, Gly tRNA; tRNA2, Met tRNA. **B-D**) The pegRNAs were designed to tag *OsMPK6* at the GAG PAM site (**B**), *OsMPK7* at the GAG PAM site (**C**), and *OsMPK10* at the TAG PAM site (**D**) through ScCas9-mediated NM-PE. **E-G**) Representative Sanger sequencing chromatograms display *in situ* FLAG epitope tagging at the *OsMPK6* site (**E**), c-Myc epitope at the *OsMPK7* site (**F**), and FLAG epitope at the *OsMPK10* site (**G**). **H**) Summary of the gene tagging frequencies of rPE16b in T0 transgenic rice lines. Rice with heterozygous, homozygous, and biallelic tag insertions is designated as He, Ho, and Bi, respectively. **I**) The stacked diagram depicts the proportions of distinct editing outcomes in alleles created by rPE16b. **J**) A schematic outlines the gene tagging mechanism facilitated by Dual-PE strategy. **K**) A comparison of gene tagging efficiency between NM-PE and Dual-PE strategies in T0 transgenic rice plants.

It’s been well documented that PE with a pair of pegRNAs is capable of enhancing the editing efficiency in plants (Lin et al. 2021). The ScCas9n-based prime editor, with its flexibility in selecting NNG PAM sites across target regions, could be particularly advantageous for this strategy. To evaluate whether paired pegRNAs could further improve the efficiency of *in situ* epitope tagging, we engineered the ScCas9 nuclease into its nickase version, ScCas9(H849A), resulting in rPE16a (Figure 3A). We designed paired pegRNAs in trans (termed Dual-PE), each bearing identical epitope tags for the forward and reverse strains, and assembled them into tRNA-pegRNA-HDV repeats for the individual tagging of *OsMPK6*, *OsMPK7*, and *OsMPK10* (Figure 3A, J; Supplementary Figure S6A-C). To be mentioned, the two cleavage sites of the pegRNAs pairs in the three target genes were 24 to 37-nt apart. Forty-eight independent transgenic lines were chosen to be genotyped for each gene. Among them, we found that 8 lines (16.67%) exhibited precise c-Myc tagging at the *OsMPK7* site. Notably, all positive plants were heterozygous, and no other editing events were observed (Supplementary Figure S6E, G). As of *OsMPK10*, only 2 lines (4.17%) positive for precise FLAG tagging were isolated, of which 1 line was heterozygous while the other carried biallelic edits accompanied by byproducts. Meanwhile, 5 lines were characterized by various imprecise tag insertions at either pegRNA-binding site, implying only one pegRNA, rather than both, functioned in this case (Supplementary Figure S6F, G). Regarding *OsMPK6*, no edits of any type was detected (Supplementary Figure S6D, G). Collectively, these findings indicate that the insertion efficiencies of ScCas9n-mediated dual-PE were significantly lower (more than 3.70-fold) than that of ScCas9-mediated NM-PE at the tested sites (Figure 3K). We assume that it might resulted from the ineffective recognition of non-canonical PAM sequences by ScCas9-derived prime editor, potentially impacting its ability to simultaneously cleavage two distinct sites, an essential aspect of the dual-pegRNA approach (Lin et al. 2021).

Combining all the data, we conclude that NM-PE is compatible with other engineered Cas9 variants and CRISPR/Cas systems, thereby significantly expanding its applicability for *in*-*situ* epitope tagging across the rice genome. Notably, ScCas9-mediated NM-PE enables the more extensive tagging of rice genes compared to SpCas9.

### Dual gene tagging in rice by NM-PE

Dual gene tagging is of great value in investigating protein interactions, gene regulation, signal transduction pathways, *etc*., within complex biological processes in plants. Therefore, we investigated whether NM-PE could be employed for dual gene tagging in rice plants (Figure 4A). Two pegRNAs were designed, assembled into tRNA-pegRNA-HDV array, and deliver into rice cell together with rPE14b, aiming to simultaneously insert the c-Myc and HA tags immediately upstream of the stop codons of the endogenous *OsMPK1* and *OsMPK13* genes, respectively. Out of 48 independent transgenic lines obtained, 11 and 13 lines were identified with accurate *in situ* epitope tagging at the *OsMPK1* and *OsMPK13* sites, respectively, of which 6 lines (12.50%) harbored the desired *OsMPK1-c-Myc* and *OsMPK13-HA* genes simultaneously (Figure 4B-E). The immunoblotting analysis revealed the presence of 45 kDa of *OsMPK1* and 58 kDa of *OsMPK13* in T0 plants using c-Myc and HA antibodies, respectively, indicating that both endogenous *OsMPK1* and *OsMPK13* genes were precisely tagged and expressed (Figure 4F). Alternatively, like single gene tagging, a significant number of plants carried byproducts, primarily consisting of Indels and inaccurate insertions (Supplementary Figure S7). Nevertheless, transgene-free plants homozygous for both single and dual gene tagging are expected to be isolated from the offsprings of certain T0 plants (line #1, #4, #28, *etc*.) after self-pollination. These data demonstrate the feasibility of using the NM-PE strategy for single, dual or even multiple gene tagging in rice.

**Figure 4.**
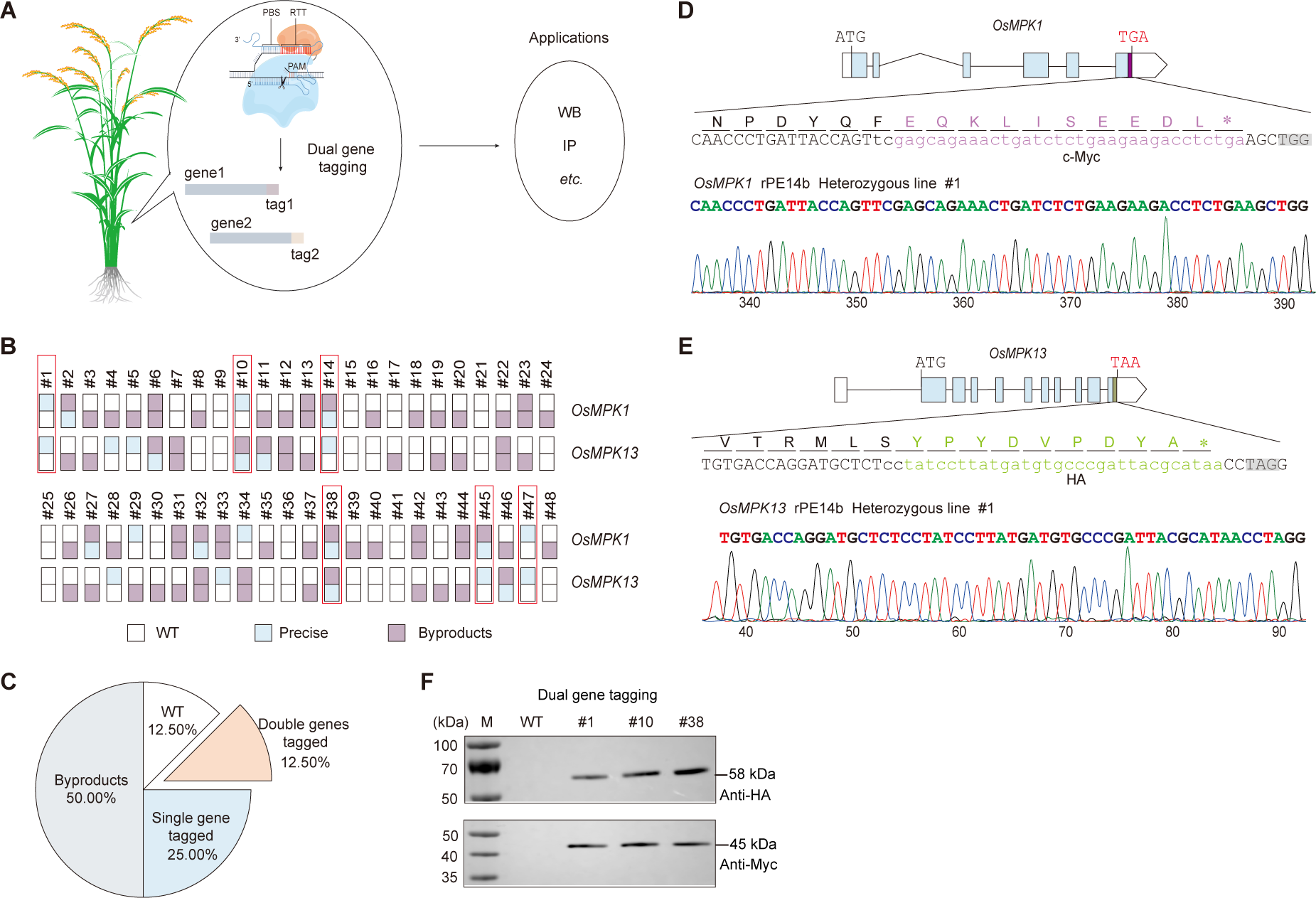
Dual gene tagging in rice by NM-PE. **A**) A schematic illustrates the application of dual gene tagging in plant studies. **B**) Genotyping of dual gene tagging in T0 transgenic rice lines were conducted, with plants bearing dual gene tags highlighted by red frames. **C**) A pie chart quantifies the distribution of various gene tagging types created by NM-PE-mediated dual gene tagging in T0 transgenic rice lines (n = 48). **D-E**) Representative Sanger sequencing chromatograms of dual gene tagging created by NM-PE. *OsMPK1* was tagged with the c-Myc epitope (**D**) and *OsMPK13* was tagged with the HA epitope (**E**). **F**) Immunoblotting analysis revealed the presence of 45 kDa of *OsMPK1* and 58 kDa of *OsMPK13* in T0 plants using c-Myc and HA antibodies, respectively. Total protein extracts from non-transgenic rice plants served as negative controls.

## Discussion

Considerable efforts have been devoted to advancing *in situ* epitope tagging technologies. However, its broad application in plants remains challenging. In this study, the derivative NM-PE strategy, utilizing CRISPR/Cas nuclease and the MMEJ repair mechanism, has been developed. We demonstrate that this strategy, compatible with SpCas9, SpRY, and ScCas9, facilitates precise insertion of diverse epitope tags immediately preceding the stop codon of endogenous rice genes, achieving a tagging efficiency of up to 70.83% in rice plants. Moreover, it allows for efficient multiplex gene tagging with tRNA-pegRNA-HDV array.

Along with the development of genome editing technologies, a variety of nucleases, specially SpCas9, have been successfully employed in the insertion and replacement manipulation of DNA fragment in various organisms, utilizing exogenous DNA/RNA donors through either NHEJ or HDR pathway. By contrast, NM-PE employs a 3’ DNA flap containing the desired epitope tag sequence, replicated from pegRNAs through reverse transcription by M-MLV, thus achieving efficient endogenous gene tagging. This approach resembles the utilization of DNA-binding proteins fused with CRISPR/Cas system to enrich donor DNA at DSB sites, thereby increasing the efficiency of nuclease-based fragment manipulation (Aird et al. 2018; Wang et al. 2021a). However, NM-PE eliminates the need for exogenous donor templates, rendering it more cost-effective through *Agrobacterium*-mediated transformation, particularly for large-scale endogenous gene tagging in plants. Nevertheless, NM-PE, like the conventional PE, is constrained by the reverse transcriptase’s activity with respect to the length of the insertion fragment. Next, we intend to investigate longer inserts using alternative reverse transcriptases with enhanced processing capabilities (Doman et al. 2023).

In agree with previous reports on PE-mediated endogenous gene tagging in plants, our extensively optimized high-efficiency rPE14a, as studied here, achieved a gene tagging efficiency of 22.82% in transgenic rice plants. However, NM-PE exhibited a remarkable improvement of over 2.50-fold in gene tagging compared to rPE14a-mediated PE at the tested sites. It is highly likely attributed to NM-PE’s robust ability to induce DSBs at the target site, thereby facilitating efficient repair through the MMEJ pathway downstream. This trend has also been observed in animal cells when employing nuclease-mediated derivative PE for base substitutions, small fragment insertions, and deletions (Adikusuma et al., 2021; Jiang et al., 2022a; Peterka et al., 2022; Tao et al., 2022; Li et al., 2023b). However, unlike in animal cells, our results in plant cells show that even without homology arms, the 3’ Flap can undergo MMEJ repair by seeking 2-5-nt microhomology sequences in the flanking regions (Supplementary Figure S3), whereas animal cells tend to prefer NHEJ repair. This phenomenon may be attributable to inherent differences in repair mechanisms between plant and animal cells. Although NN-PE can insert complete tags into the genome by seeking flanking microhomology sequences, this spontaneous MMEJ repair may result in deletions, potentially compromising the functionality of the tagged protein. However, NM-PE does not produce such deletions, thereby suggesting that the NM-PE strategy provides a distinct advantage for gene tagging in plants. Furthermore, unlike the GRAND editing approach, which employs two pegRNAs, NM-PE present a more simplified pegRNA design, wherein a shorter RTT ensures precise gene tagging, high efficiency, and eliminates the risk of nucleotide loss of target region-a common occurrence in GRAND editing. Particularly in the case of the ScCas9-based system, NM-PE optimally enables the insertion of epitope tags immediately upstream of the stop codon of the target genes, thereby preserving the integrity of the coding sequences (adding at most one extra amino acid) and minimizing the impact on the functionality of the tagged protein. Moreover, NM-PE confers a significant advantage in facilitating multiplex gene tagging with a simple tRNA-pegRNA-HDV repeat assembly, enabling researchers to explore the synchronized expression and interaction dynamics of multiple genes within the native context of plant cells. It is worth noting that, in comparison to the nickase-based PE, nuclease-based NM-PE leads to a noticeable increase in byproducts, particularly Indels. Nevertheless, accurate gene tagging events typically occur independently of Indel mutations on different chromosomes in plant cell, allowing for the isolation of transgene-free, homozygous gene-tagged individuals in the next generation population. Anyhow, we examined potential off-target sites predicted by the online tool CRISPR-GE in this study, and no editing was observed at these potential off-target sites (Supplementary Figure S8).

In addition to SpCas9, the PAM-flexible SpRY and ScCas9 significantly expand the targeting scope of NM-PE in rice. To investigate the potential application of NM-PE for genome-wide *in situ* epitope tagging, we searched for all potential target sites of NM-PE in 48,494 transcripts within *Oryza sativa* Kitaake genome V3 from Phytozome (https://data.jgi.doe.gov/refine-download/phytozome). We discovered that SpCas9-mediated NM-PE can target 72.70% of transcripts at the N-terminus and 41.31% at the C-terminus, accounting for 84.06% of all transcripts (Supplementary Figure S9). Meanwhile, ScCas9-mediated NM-PE approaches nearly 100% coverage of all transcripts at both termini. Moreover, the HiBiT tag (11 amino acids), generates bright luminescence through high-affinity complementation with an 18 kDa subunit derived from NanoLuc (LgBiT), enabling the quantitative assessment of dynamic processes associated with the regulated expression and covalent modifications of endogenous proteins (Schwinn et al. 2018; Tian et al. 2024). Based on these findings, we believe that the combined use of the HiBiT tag and NM-PE in the future will greatly accelerate the progress of the Rice Protein Tagging Project (Lu et al. 2020a).

In conclusion, we have successfully developed a straightforward and highly efficient derivative prime editing strategy, termed NM-PE, for precise epitope tagging of endogenous rice genes. This method allows for convenient insertion of any epitope tag sequence immediately prior to the stop codon, enabling single as well as dual gene tagging throughout the genome using a straightforward pegRNA design and assembly process. Our comprehensive research provides alternative perspectives on the advancement of *in situ* epitope tagging technologies in other plants and will undoubtedly accelerates the Rice Protein Tagging Project, gene function study and genetic improvement in the future.

## Materials and Methods

### Rice cultivars and growth conditions

Rice cultivars Kitaake (*Oryza sativa* L. ssp. *Geng*) was utilized for this study and maintained in our laboratory. In the paddy field, rice plants were grown naturally throughout the regular rice growing season, and immature seeds were collected for rice transformation.

### Plasmid construction

To construct PE vectors, the coding sequences of the Moloney murine leukemia virus reverse transcriptase pentamutant without the RNase H domain (M-MLV RT-ΔRNase H/D200N/L603W/T330P/ T306K/W313F) were codon-optimized for expression in rice, named M-MLV (ΔRNase H) (Supplementary Table S1), and synthesized (Tsingke Biotech, China). The SpCas9 in pUC57:SpCas9 was reverted to SpCas9 (R221K/N394K/H840A) by the Mut Express II Fast Mutagenesis Kit V2 (C214, Vazyme, China) (Zhou et al. 2014), resulting in pUC57:SpCas9 (R221K/N394K/H840A). Next, the synthesized fragment in pUC57:M-MLV (ΔRNase H) was used as a template to amplify fragment 1, with the primer pairs rPE14-F1/rPE14-R1. The full-length *SpCas9* (R221K/N394K/H840A) gene was PCR amplified using pUC57:SpCas9 (R221K/N394K/H840A) as the template to generate fragment 2 by the primer pair rPE14-F2/rPE14-R2. Fragments 1 and 2 were ligated by the ClonExpress II One-Step Cloning Kit (C112, Vazyme, China), resulting in pUC57:rPE14a. The chimeric gene in pUC57:rPE14a was used to substitute the *SpRY* gene between the maize ubiquitin 1 promoter and the NOS terminator in pUbi:SpRY through *Bam*H I/*Spe* I digestion and DNA ligation (Xu et al. 2021), resulting in the binary vector pUbi:rPE14a.

To develop derivative PE systems that leverage nuclease-mediated PE. With the same strategy, the SpCas9 (R221K/N394K/H840A) in pUC57:rPE14a was reverted to its wild-type form, resulting in pUC57:rPE14b. To expand the range of targetable genes in rice and explore whether the NM-PE strategy are compatible with other SpCas9 variants (SpRY) and an ortholog (ScCas9) that recognize relaxed PAMs (Wang et al. 2020; Xu et al. 2021). The primer pair rPE15-F1/rPE15-R1 was used to amplify the full-length *SpRY* gene and pUbi:SpRY as the template to construct pUC57:rPE15b (Xu et al. 2021). The primer pair rPE16-F1/rPE16-R1 was used to amplify the full-length *ScCas9* gene and pUbi:ScCas9 as the template to construct pUC57:rPE16b (Wang et al. 2020). For the construction of the binary vector of ScCas9 (H849A) nickase, a point mutation of CAT to GCT (H849A) was first introduced into ScCas9 by PCR amplification of the whole plasmid pUC57:rPE16b with the primer pair rPE16-F2/rPE16-R2. The amplicon was self-ligated and confirmed by sequencing, resulting in pUC57:rPE16a. Finally, pUbi:rPE14b, pUbi:rPE15b, pUbi:rPE16a, and pUbi:rPE16b were constructed using the same strategy as previously described.

To construct an entry vector for pegRNA expression, the *Bam*H I/*Eco*R I digestion fragment of pENTR4:sgRNA4 was replaced with a synthetic fragment of 35S-CmYLCV-U6-tRNA-*Bsa* I-*Bsa* I-evopreQ1-HDV-polyT-HSPt (Supplementary Table S1), resulting in pENTR4:sgRNA41 (Yan et al. 2021). The PBS with a melting temperature of 30°C was designed using the plant PegDesigner version 1.0 (http://www.plantgenomeediting.net). An 8-nt linker was predicted by the pegLIT tool (http://peglit.liugroup.us) to connect the structured RNA motif sequence of evopreQ1. Then, the fragments comprising the protospacer-sgRNA-RTT-PBS-linker (Supplementary Table S1) were chemically synthesized and digested with terminal restriction sites (*Bsa* I or *BtgZ* I). Subsequently, they were inserted into the *Bsa* I site of pENTR4:sgRNA41 by *Bsa* I digestion and DNA ligation. Finally, the pegRNA expression cassettes were inserted into different binary vectors through the LR reaction and transformed into *Agrobacterium* EHA105 for rice callus transformation to tag the *OsMPKs* (Zhou et al. 2014).

All the PCR fragments for plasmid construction were amplified with TransStart FastPfu PCR SuperMix (AS221, TransGen Biotech, China), and the identities of all recombined plasmids were confirmed by Sanger sequencing. Table S2 (Supplementary) contains a list of all primers utilized in this investigation.

### *Agrobacterium*-mediated rice transformation

The relevant plasmids were introduced into the *A. tumefaciens* strain EHA105 through electroporation. Transgenic plants were generated through *Agrobacterium*-mediated transformation with immature rice seeds as mentioned previously (Wang et al. 2022b).

### Genomic DNA extraction and genotyping

Genomic DNA was extracted from leaves of T0 transgenic plants using the hexadecyltrimethylammonium bromide (CTAB) method (Porebski et al. 1997). The target region of each endogenous gene was PCR amplified using 2x Rapid Taq Master Mix DNA polymerase (P222, Vazyme, China) with gene-specific primers listed in Supplementary Table 3. Equal amount of PCR products with different barcodes were mixed and subjected to sequencing, generating approximately one gigabyte of data using the Illumina NextSeq platform with a paired-end 150 bp (PE-150) strategy. The precisely tagged plants were randomly chosen and further validated by Sanger sequencing.

### Prediction and investigation of the sgRNA-dependent off-target sites

To evaluate the specificity of the NM-PE systems, the potential off-target sites were predicted by the online tool CRISPR-GE (https://skl.scau.edu.cn) (Xie et al. 2017) and BLAST in the Rice Annotation Project Database (https://rapdb.dna.affrc.go.jp). The potential off-target regions were PCR amplified with specific primers, and the PCR products were subjected to deep sequencing using the Illumina NextSeq platform.

### Total protein extraction and immunoblot analysis

The rice leaves of T0 plants were individually treated with abscisic acid (ABA, 2.46 μg/mL) and chitin (0.1 μg/mL) for 1 h at room temperature (Shi et al. 2014). All leaf tissues from the same plant were harvested together and ground into fine power with liquid nitrogen. Total protein was isolated from 0.1 g of samples using an extraction buffer composed of 100 mM Tricine, 10 mM KCl, 1 mM MgCl2, 1 mM EDTA, 10% sucrose, 2% Triton-X100, 1 mM DTT, and a 1×protease inhibitor cocktail (RM02916, ABclonal Tech, China). After extraction, the samples were centrifugated at 12,000 rpm for 10 min at 4℃, and 8 μl of each extract with loading dye was resolved on an 8% polyacrylamide SDS-PAGE gel and electroblotted onto PVDF membranes at 22 V for 40 min. The membranes were blocked with 5% skim milk dissolved in TBST (20 mM Tris-HCl, pH 7.6, 150 mM NaCl, 0.1% v/v Tween-20). Anti-HA (AE008, ABclonal Tech, China) and anti-Myc (AE010, ABclonal Tech, China) antibodies diluted 1:5,000 in TBST with 5% skim milk were used. After primary antibody incubation, membranes were washed three times (5 min each) with TBST, then incubated with a goat anti-mouse IgG HRP-linked secondary antibody (AS003, ABclonal, China, 1:5,000 dilution) in TBST containing 5% skim milk for 1 h at room temperature. Subsequently, the membranes were washed three times (10 min each) in TBST. Signals were detected using the Super ECL Detection Reagent (36208ES60, Yeasen, China) and a touch imager (e-BLOT, China).

### Accession numbers

Sequence data from this article can be found in the Rice Annotation Project Database (RAP-DB) under the following accession numbers: *OsACC,* LOC_Os05g22940; *OsMPK1*, LOC_Os06g06090; *OsMPK3*, LOC_Os02g05480; *OsMPK4*, LOC_Os06g48590; *OsMPK6*, LOC_Os10g38950; *OsMPK7*, LOC_Os05g49140; *OsMPK8*, LOC_Os01g47530; *OsMPK10*, LOC_Os01g43910; *OsMPK11*, LOC_Os06g26340; and *OsMPK13*, LOC_Os02g04230.

## Acknowledgements

We thank Meixia Wang, Zhongming Zhang and Xin’ge Li for assistance with plasmid construction and genomic DNA extraction. The project was supported by the Biological Breeding-Major Projects (2023ZD04074), the National Key Research and Development Program of China (2023YFD1202905), the Nanfan special project of the Chinese Academy of Agricultural Sciences (YBXM2313), the Hainan Seed Industry Laboratory (B23CJ0208), and the Agricultural Science and Technology Innovation Program of the Chinese Academy of Agricultural Sciences.

## Author contributions

H.Z., J.H., C.S., and X.Z. designed the research; X.L., S.Z., F.Y., and B.R. conducted the experiments; C.W. and S.L. performed the bioinformatics analysis; H.Z., B.R., J.H., and X.Z. supervised the research; X.L., S.Z., H.Z., and C.S. wrote the original draft; all authors participated in discussion and revision of the manuscript.

## Supplementary Data

**Supplementary Figure S1.**
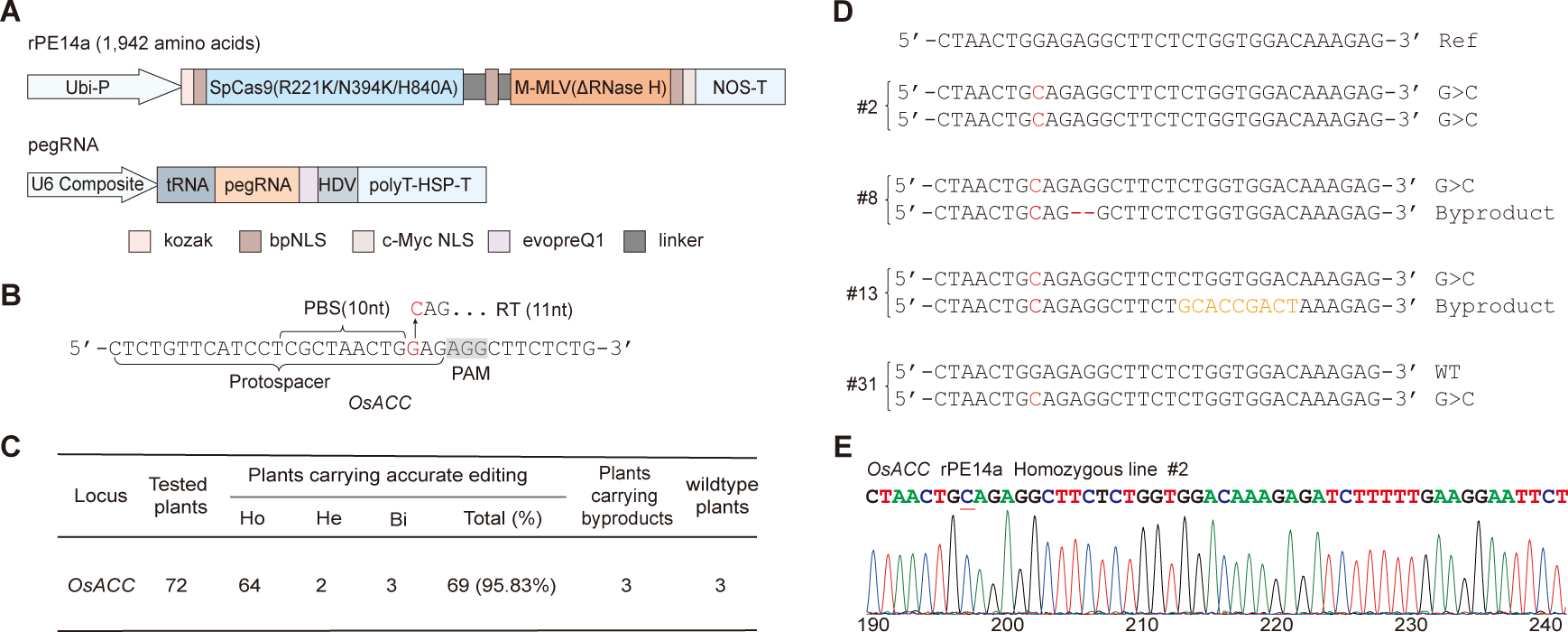
The optimized prime editor rPE14a efficiently induces G-to-C transversions at the *OsACC* site. **A)** Illustration of the rPE14a and pegRNA vectors for prime editing. Ubi-P, maize ubiquitin 1 promoter; M-MLV RT (ΔRNase H), Moloney murine leukemia virus reverse transcriptase pentamutant (D200N/L603W/T330P/T306K/W313F) without the RNase H domain; NLS, nuclear localization signal; NOS-T, terminator of nopaline synthase. The U6 composite promoter combines the CaMV 35S enhancer, the CmYLCV promoter, and a truncated U6 promoter. tRNA, Gly tRNA; HDV, HDV ribozyme; evopreQ1, a structured RNA motif of modified prequeosine1-1 riboswitch aptamer; HSP-T, AtHSP18.2 terminator. **B)** A pegRNA was designed for G-to-C transversion in *OsACC*. PBS, primer binding site; RT, reverse transcriptase; PAM, protospacer adjacent motif. **C)** Summary of the editing frequencies of rPE14a in T0 transgenic rice. Plants with precise editing exhibit at least one allele at the target site with an exact G-to-C transversions, categorized as homozygous (Ho), heterozygous (He), and biallelic (Bi) where edits and byproducts reside on different chromosomes. Plants with byproducts display at least one allele at the target site with imprecise insertions and/or Indels. **D)** Representative sequences of the rPE14a-induced *OsACC* mutations in T0 transgenic rice plants. The G-to-C transversion is highlighted in red; nucleotide deletions and sequence substitutions are marked by red dashes and orange characters, respectively. **E)** Representative Sanger sequencing chromatogram of the edited *OsACC* allele. The G-to-C transversion is underlined.

**Supplementary Figure S2.**
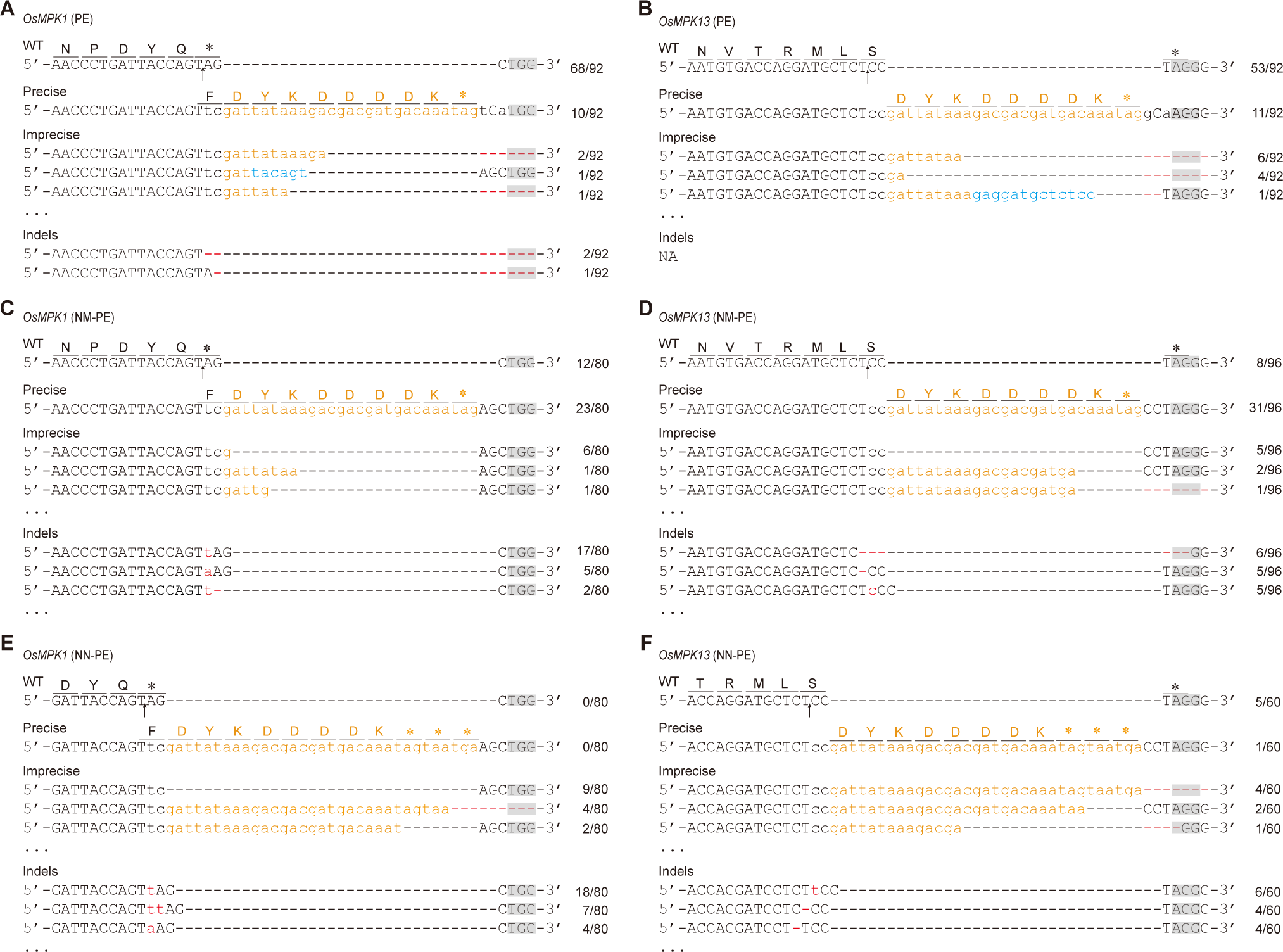
Sequencing results of the edited *OsMPK1* and *OsMPK13* in T0 transgenic rice plants generated by different PE strategies. **A-B**) Targeted gene editing of *OsMPK1* (**A**) and *OsMPK13* (**B**) by PE. **C-D**) Targeted gene editing of *OsMPK1* (**C**) and *OsMPK13* (**D**) by NM-PE. **E-F**) Targeted gene editing of *OsMPK1* (**E**) and *OsMPK13* (**F**) by NN-PE. Amino acid sequences are annotated above the corresponding DNA sequences. Insertion sites are indicated by arrows, and deletions are highlighted with red dashed lines. The insertion sequences are displayed in colored lowercase letters to distinguish between indels (red), the FLAG tag (orange), and imperfectly aligned nucleotides (blue). The protospacer adjacent motif (PAM) is labeled in gray. On the right margin, the quantity of alleles is quantified, and the three predominant gene-tagging products are depicted for each category. PE, conventional prime editing; NM-PE, nucleases/MMEJ-based prime editing; NN-PE, nucleases/NHEJ-based prime editing.

**Supplementary Figure S3.**
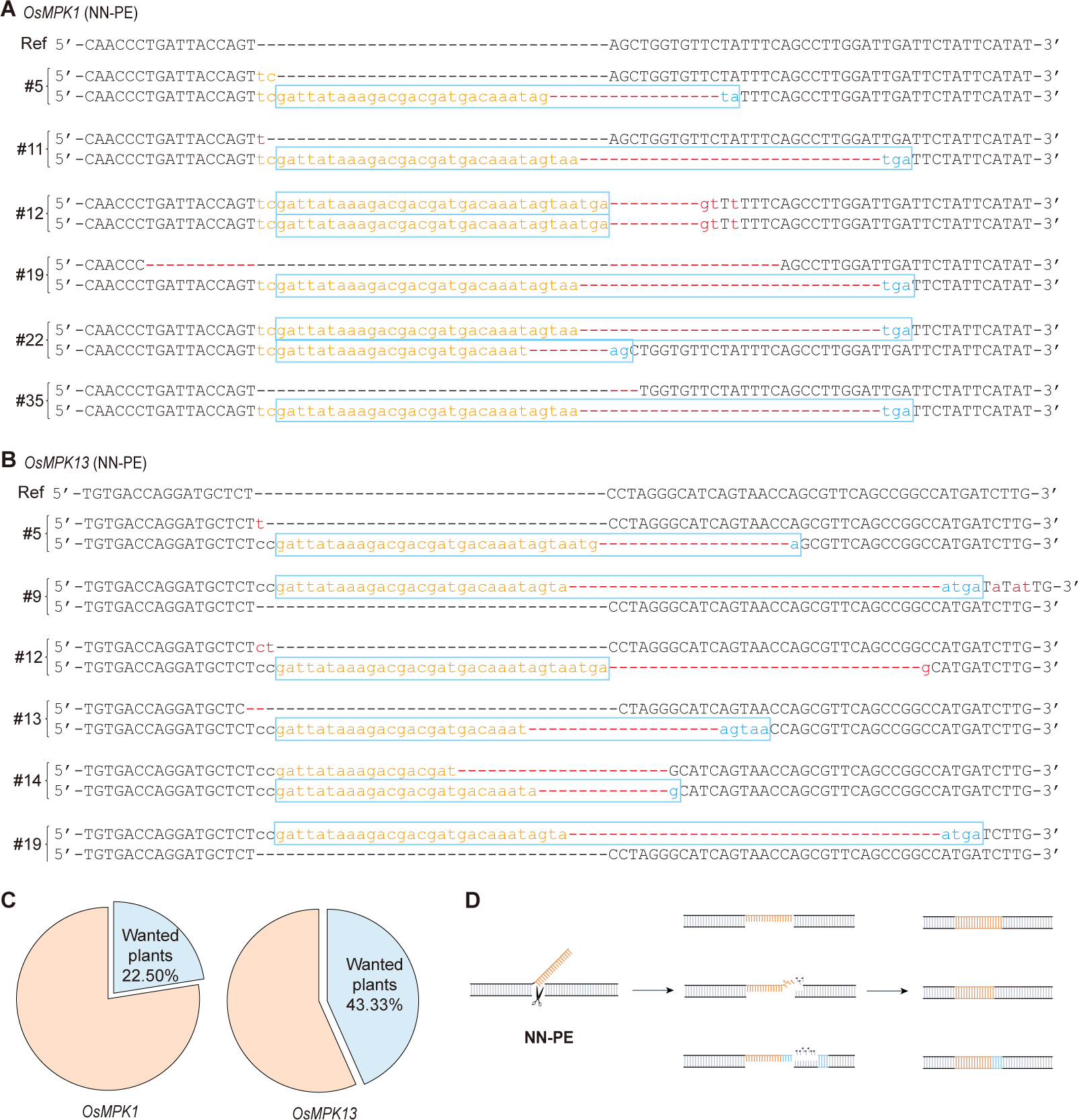
*In situ* epitope tagging of *OsMPK1* and *OsMPK13* in T0 transgenic rice plants generated via SpCas9-mediated NN-PE. **A**) Representative sequences of gene tagging created by NN-PE at the *OsMPK1* site in T0 transgenic rice plants. Six representative plants are displayed; they have incorporated the complete FLAG tag sequence and are defined as the wanted plants with epitope tagging. The complete tags are marked with blue boxes, deletions are highlighted with red dashed lines, and the insertion sequences are displayed in colored lowercase letters to distinguish between indels (red), the FLAG tag (orange), and the microhomology (blue). **B**) As for **A**), but showing the *OsMPK13* site. **C**) The proportion of the wanted plants with epitope tagging in T0 transgenic rice. **D**) A schematic illustrates the common types of gene tagging mediated by NN-PE. The tag sequences can be precisely inserted and may also produce unintended Indels, or utilize micro-homologous sequences in the downstream genome for insertion.

**Supplementary Figure S4.**
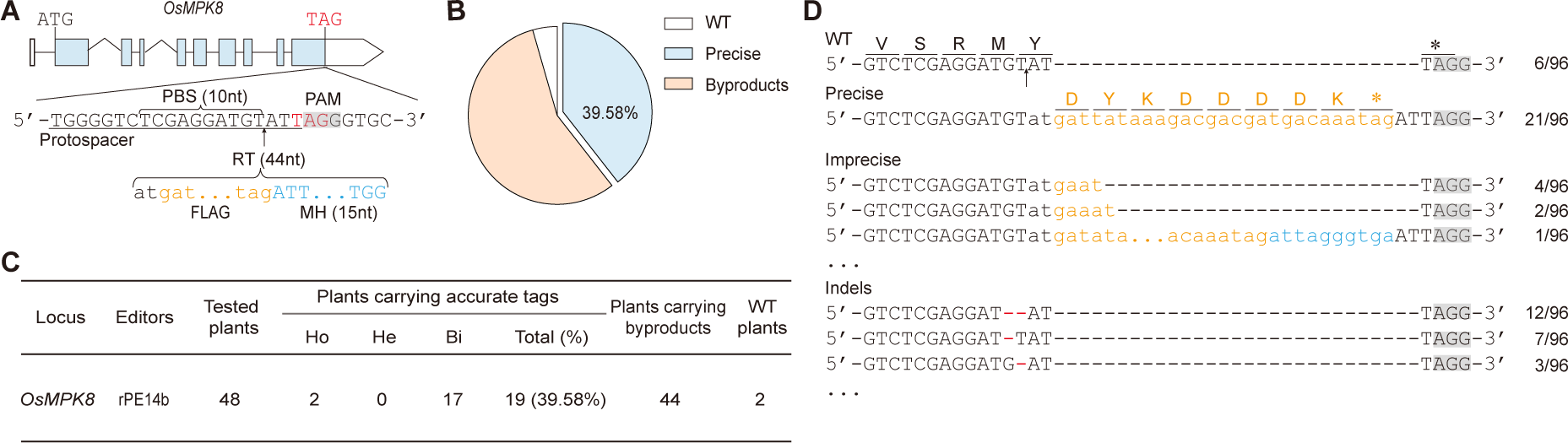
*In situ* epitope tagging of *OsMPK8* via SpCas9-mediated NM-PE. **A)** A pegRNA was designed to introduce the FLAG epitope immediately upstream of the stop codon of *OsMPK8*. **B)** The gene tagging efficiency at the *OsMPK8* locus in T0 transgenic rice (n = 48). **C)** Summary of the gene tagging frequency of SpCas9-mediated NM-PE in T0 transgenic rice lines. Plants with precise tags exhibit at least one allele at the target site with an exact insertion, categorized as homozygous (Ho), heterozygous (He), and biallelic (Bi) where edits and byproducts reside on different chromosomes. Plants with byproducts display at least one allele at the target site with imprecise insertions and/or Indels. **D)** Sequencing results of the edited *OsMPK8* in T0 transgenic rice plants. The deletions are indicated by red dashed lines. The insertion sequences are displayed in colored lowercase letters to distinguish between the FLAG tag (orange), and imperfectly aligned nucleotides (blue). In **(A)** and **(D)**, the PAM and insertion sites are highlighted in gray and marked with an arrow, respectively.

**Supplementary Figure S5.**
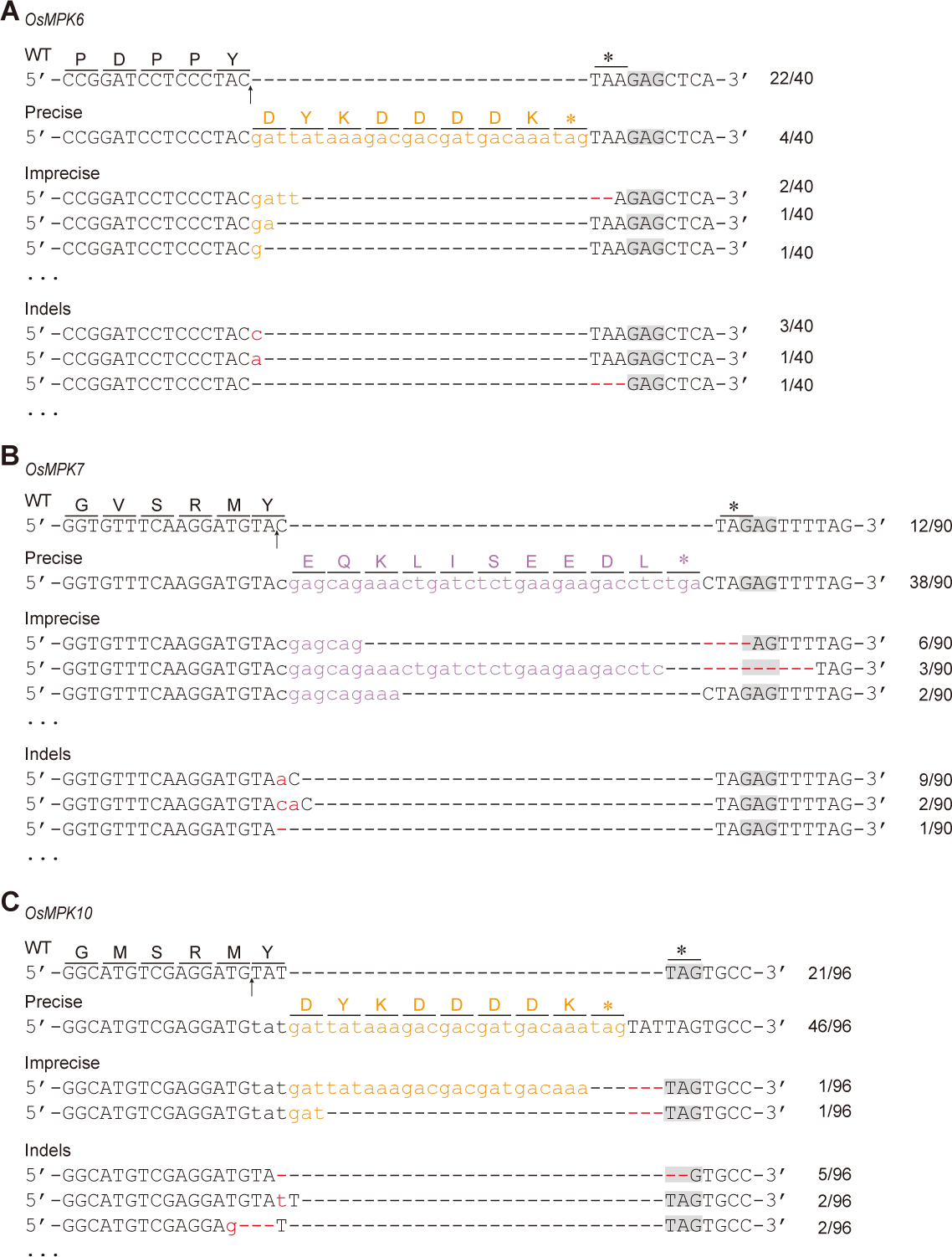
Sequencing results of the edited *OsMPKs* in T0 transgenic rice plants generated by ScCas9-mediated NM-PE. **A-C)** Targeted gene editing of *OsMPK6* **(A)**, *OsMPK7* **(B)**, and *OsMPK10* **(C)**. Insertion sites are indicated by arrows, and deletions are highlighted with red dashed lines. Inserted sequences are presented in colored lowercase letters: red for Indels, orange for the FLAG tag, and purple for the c-Myc tag. The PAM region is in gray.

**Supplementary Figure S6.**
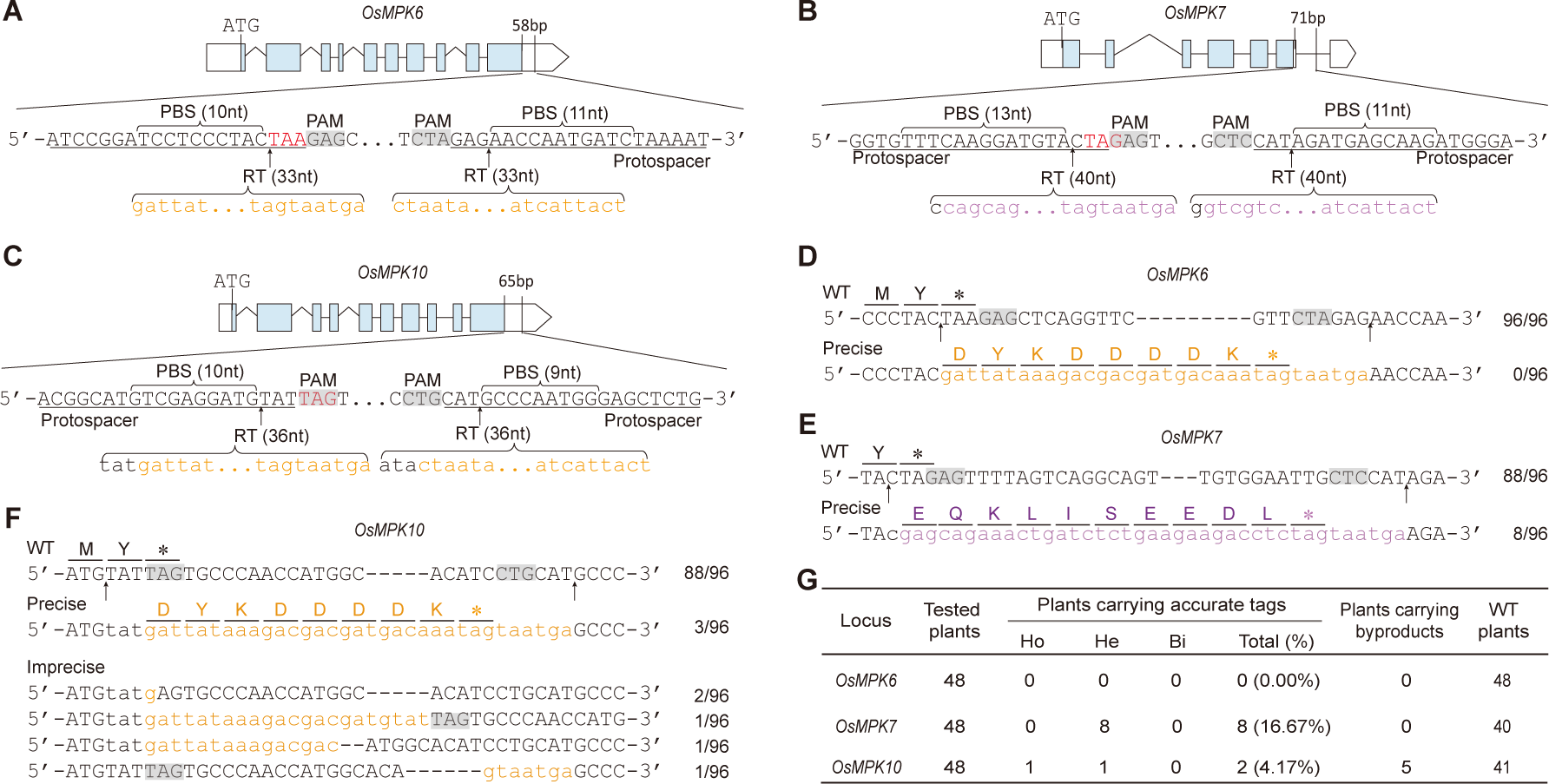
Endogenous gene tagging in rice using ScCas9n-mediated Dual-PE. **A-C)** The pair pegRNAs were designed to insert FLAG epitope tag at the *OsMPK6* site **(A)**, c-Myc epitope tag at the *OsMPK7* site **(B)**, and FLAG epitope tag at the *OsMPK10* site **(C)** through ScCas9n-mediated Dual-PE. **D-F)** Sequencing results of the edited *OsMPK6* site **(D)**, *OsMPK7* **(E)**, and *OsMPK10* **(F)** in T0 transgenic rice plants generated by ScCas9n-mediated Dual-PE. Insertion sites are indicated by arrows. Inserted sequences are displayed in colored lowercase letters: orange for the FLAG tag and purple for the c-Myc tag. **G)** Summary of the gene tagging frequencies of *OsMPKs* in T0 transgenic rice lines. Plants with precise tags exhibit at least one allele at the target site with an exact epitope-tag insertion, categorized as homozygous (Ho), heterozygous (He), and biallelic (Bi) where edits and byproducts reside on different chromosomes. Plants with byproducts display at least one allele at the target site with imprecise insertions and/or Indels.

**Supplementary Figure S7.**
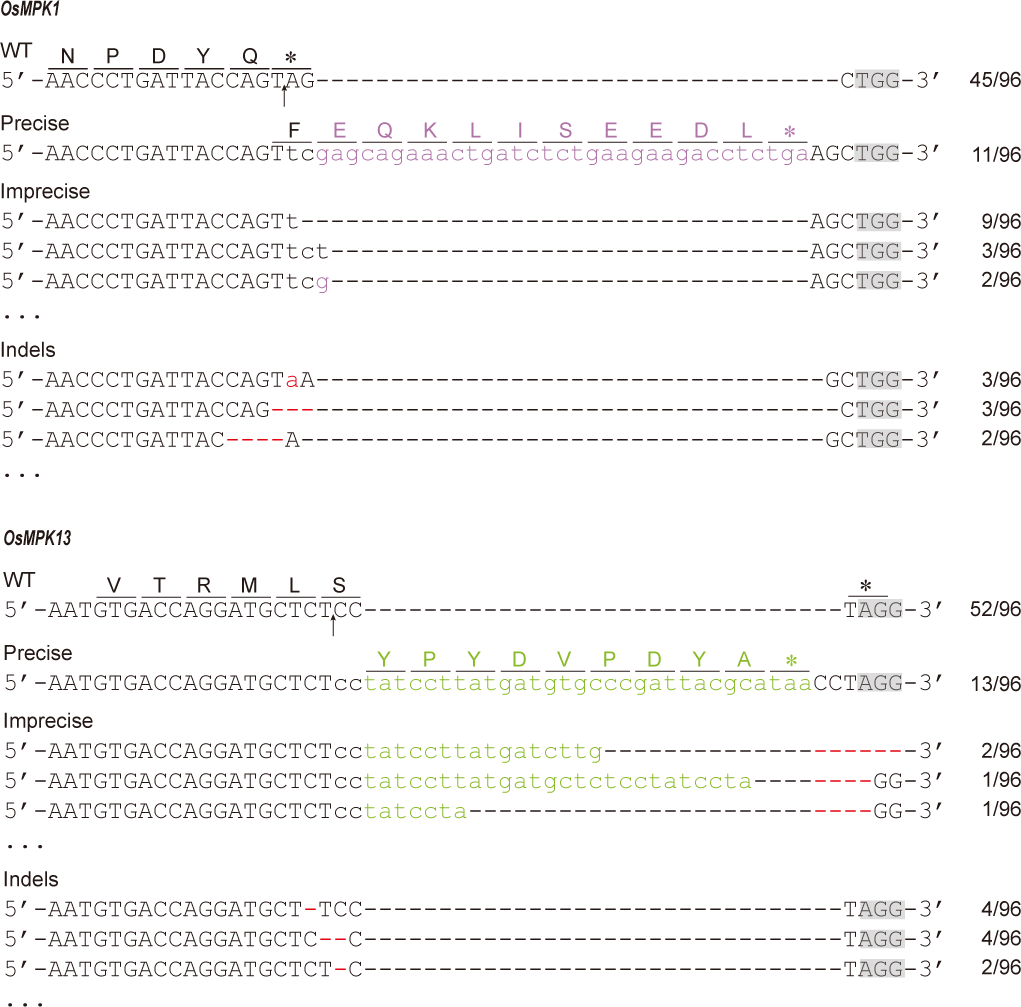
Sequencing results of the edited *OsMPK1* and *OsMPK13* in dual gene tagging in T0 transgenic plants. Insertion sites are indicated by arrows, while deletions are delineated with red dashed lines. The inserted sequences are presented in colored lowercase letters: red for Indels, purple for the c-Myc epitope, and chartreuse for the HA epitope. The PAM sequence is in gray.

**Supplementary Figure S8.**
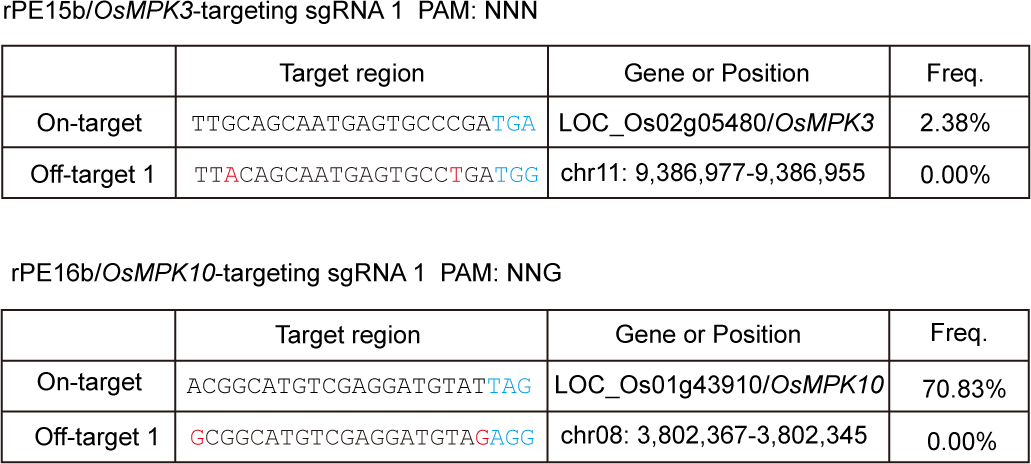
Off-target analysis of endogenous gene tagging created by NM-PE in rice. Summary of mutation frequencies in the potential off-target sites in the rice genome for nuclease in this study. The PAM sequences and the mismatches in the sgRNA sequences are highlighted in blue and red, respectively.

**Supplementary Figure S9.**
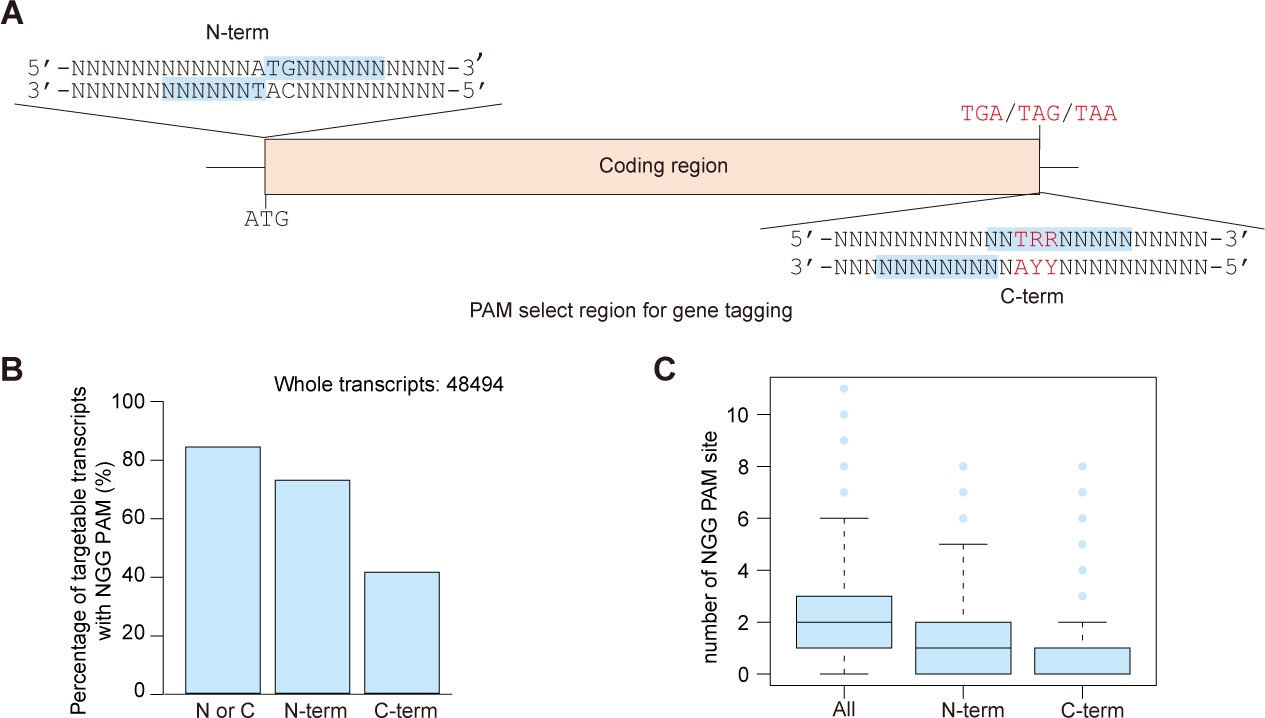
Analysis of NGG-targetable genes in the rice genome. **A)** Diagrammatic representation of the PAM selection region for *in situ* epitope tagging, with the spectrum of options denoted by blue squares. N=A, T, G, C; R=A, G. **B)** Percentage of targetable transcripts with NGG PAM in the rice genome. **C)** Distribution statistics of the number of target sites available for gene tagging with NGG PAM in each transcript in rice.

**Supplementary Table S1.**
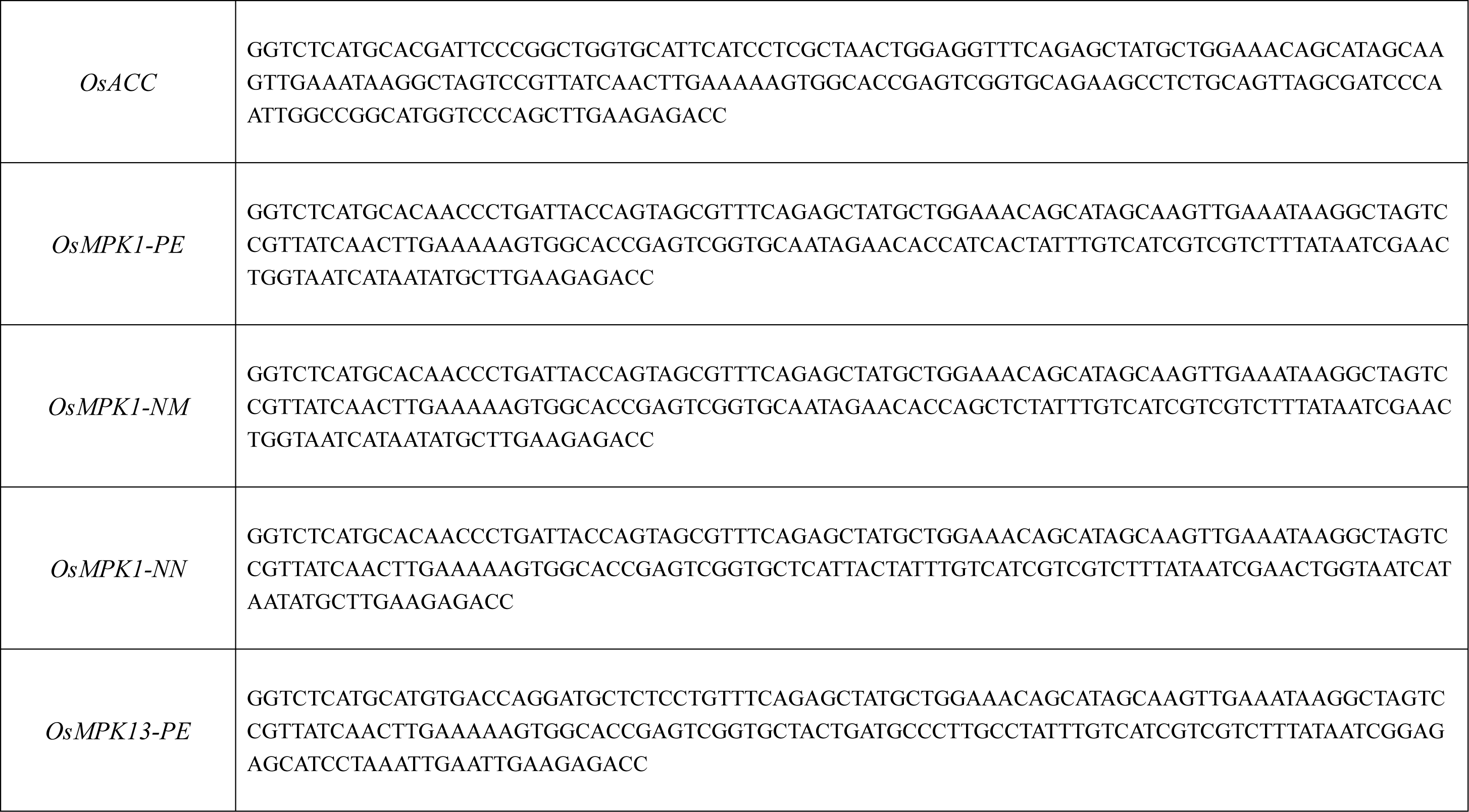
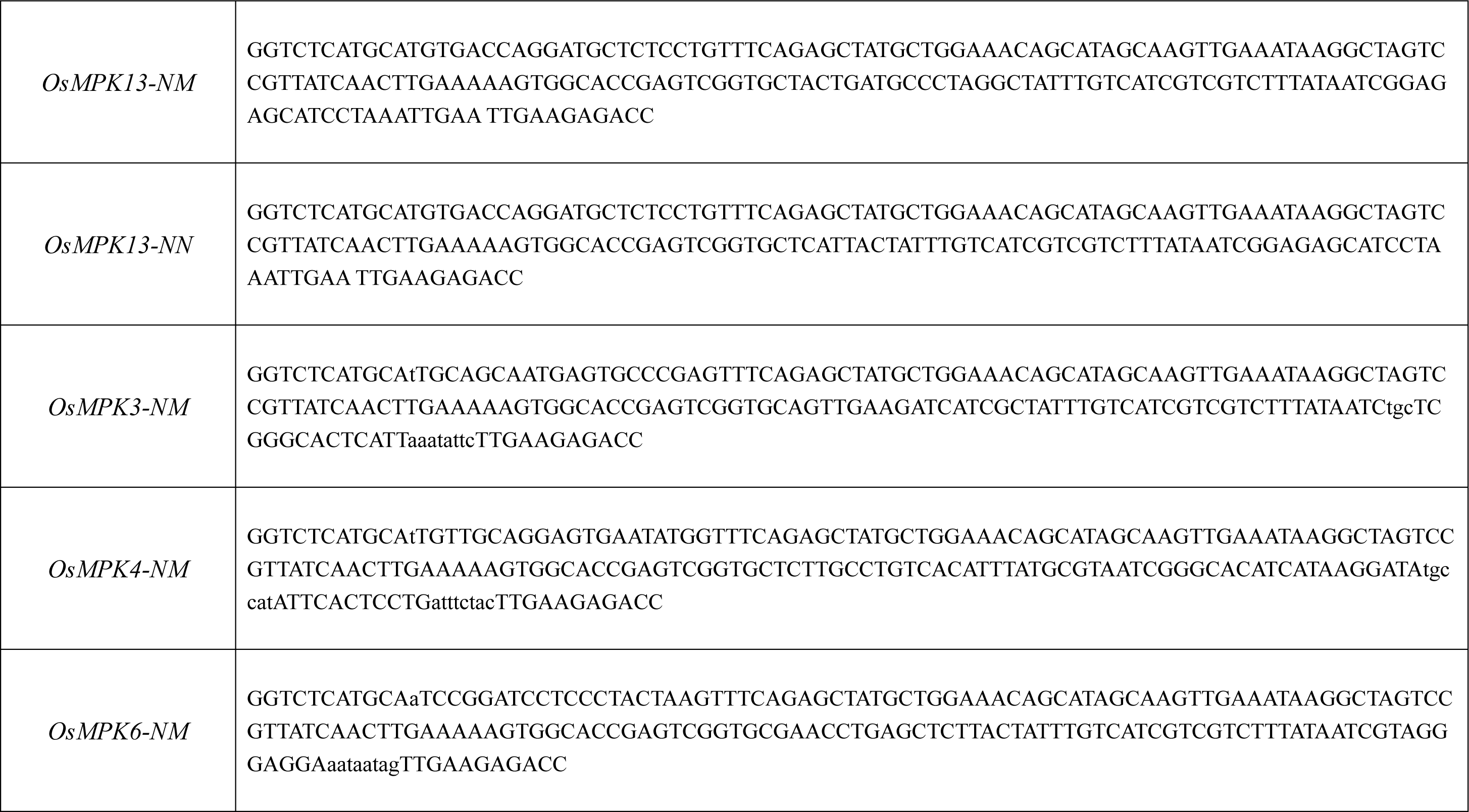
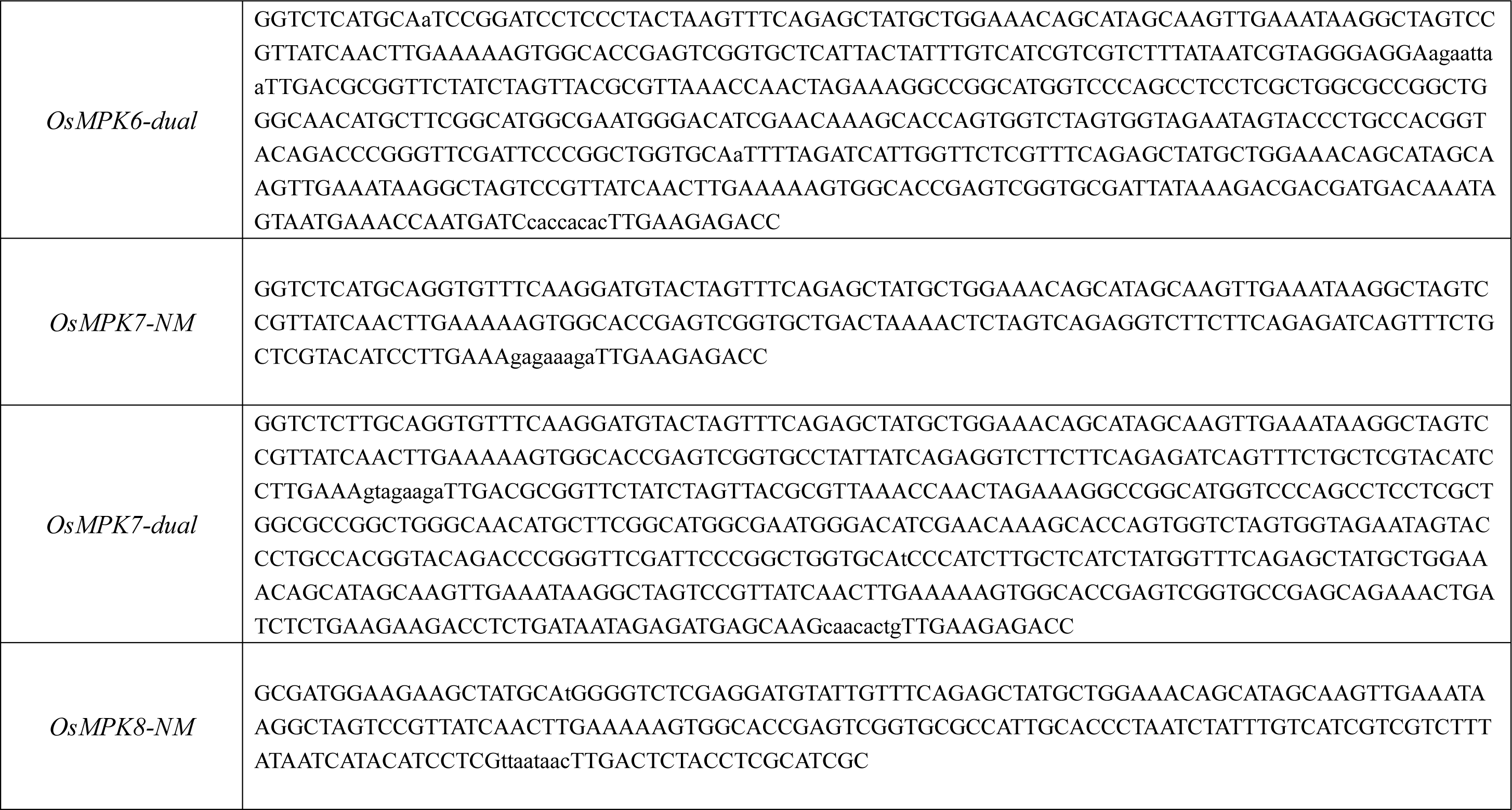
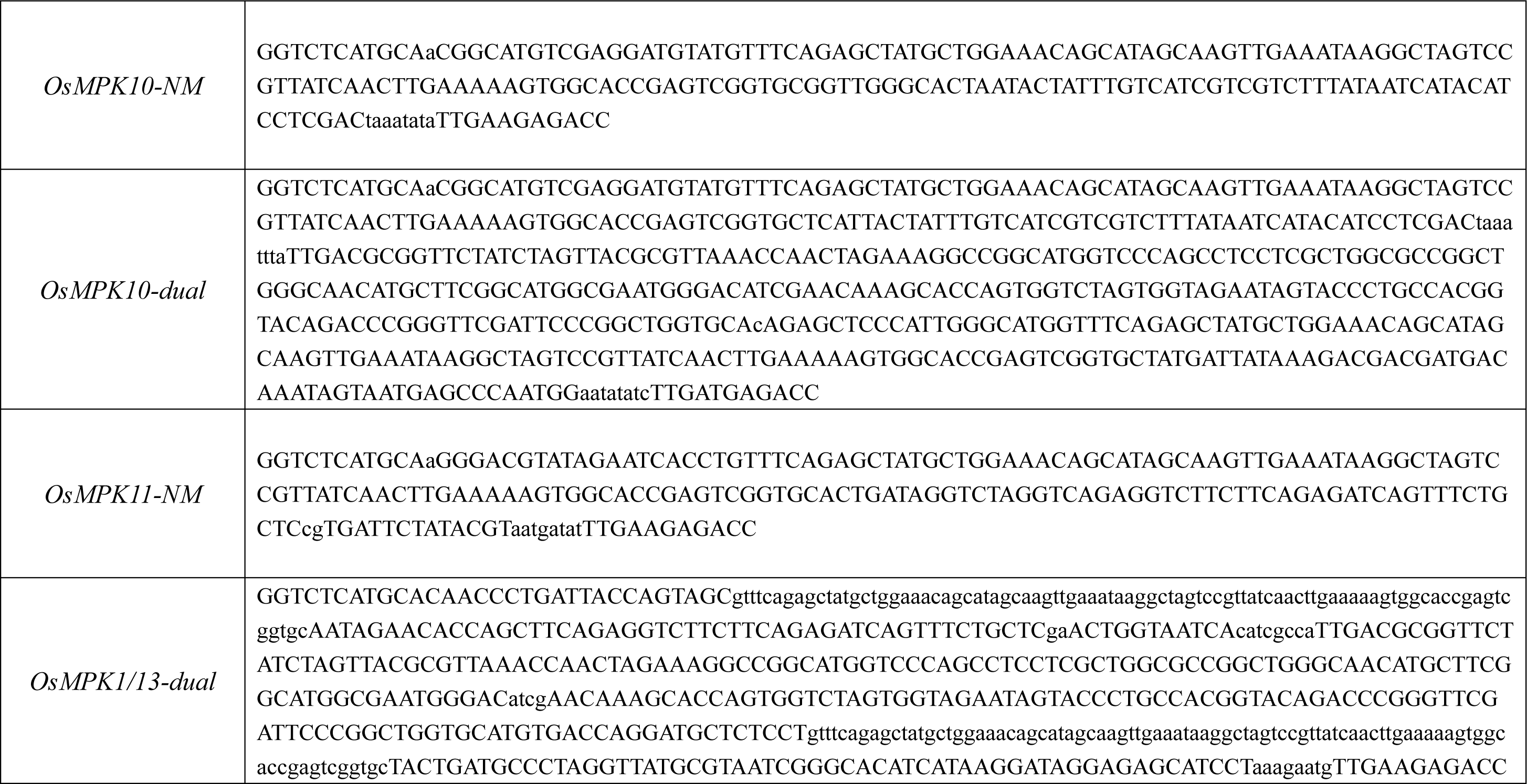
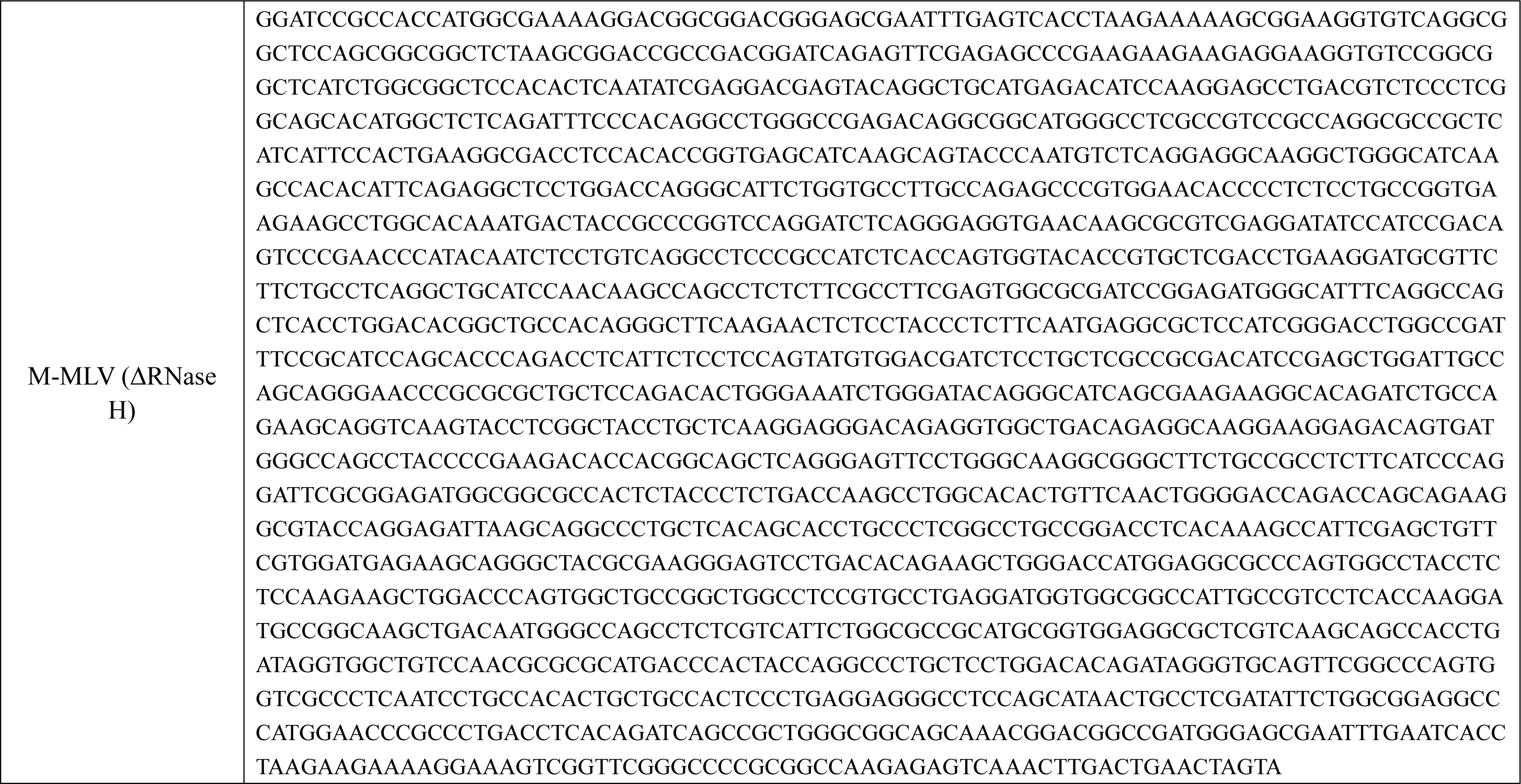
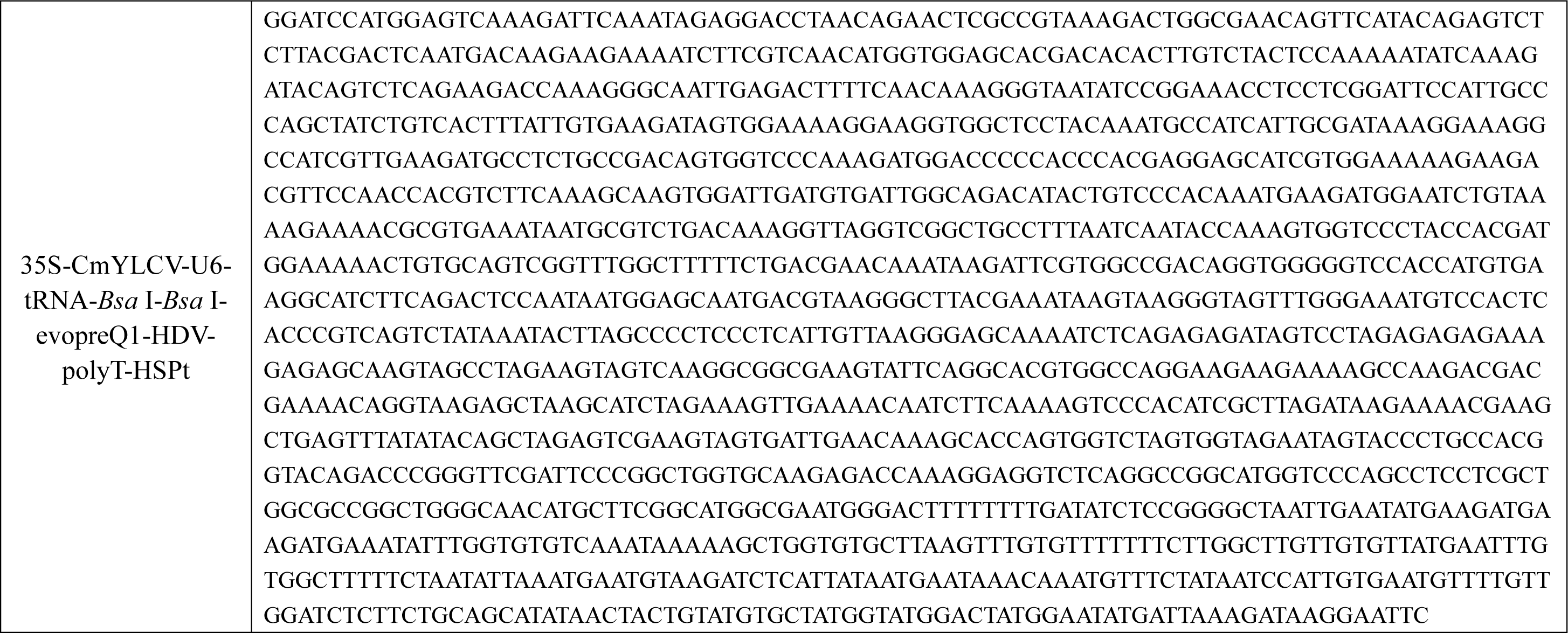
Nucleotide sequences used in this study.

**Supplementary Table S2.**
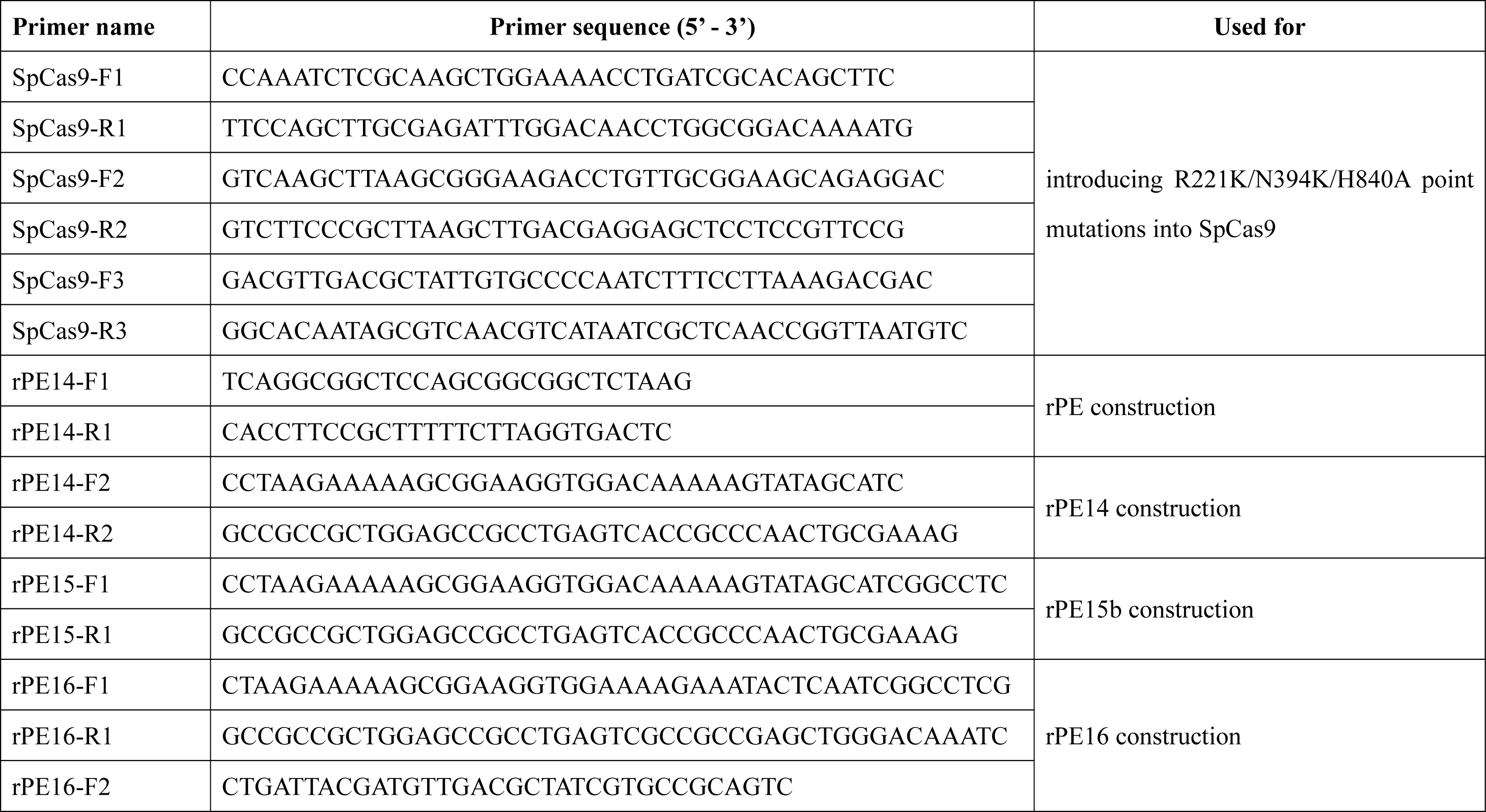
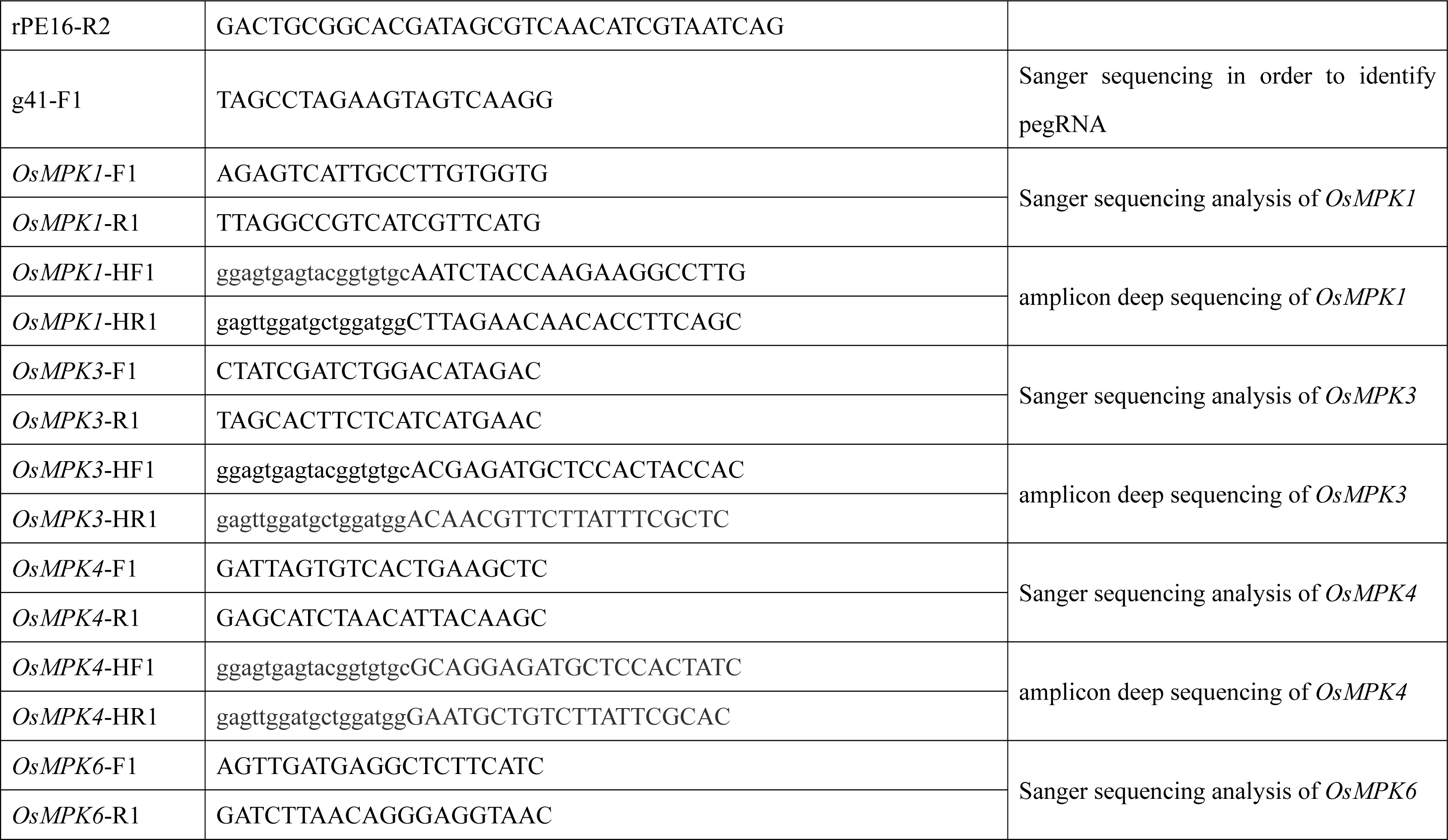
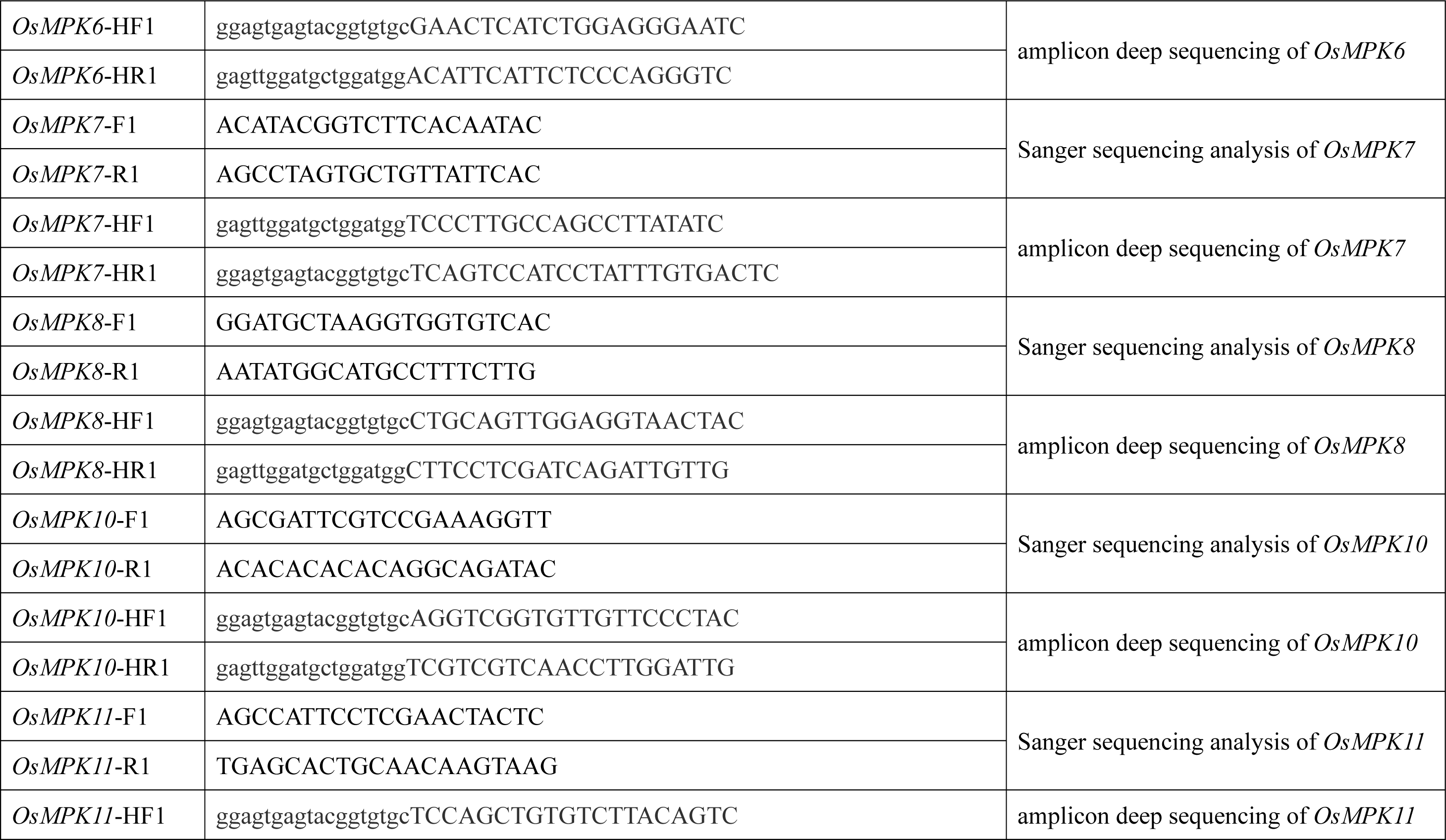
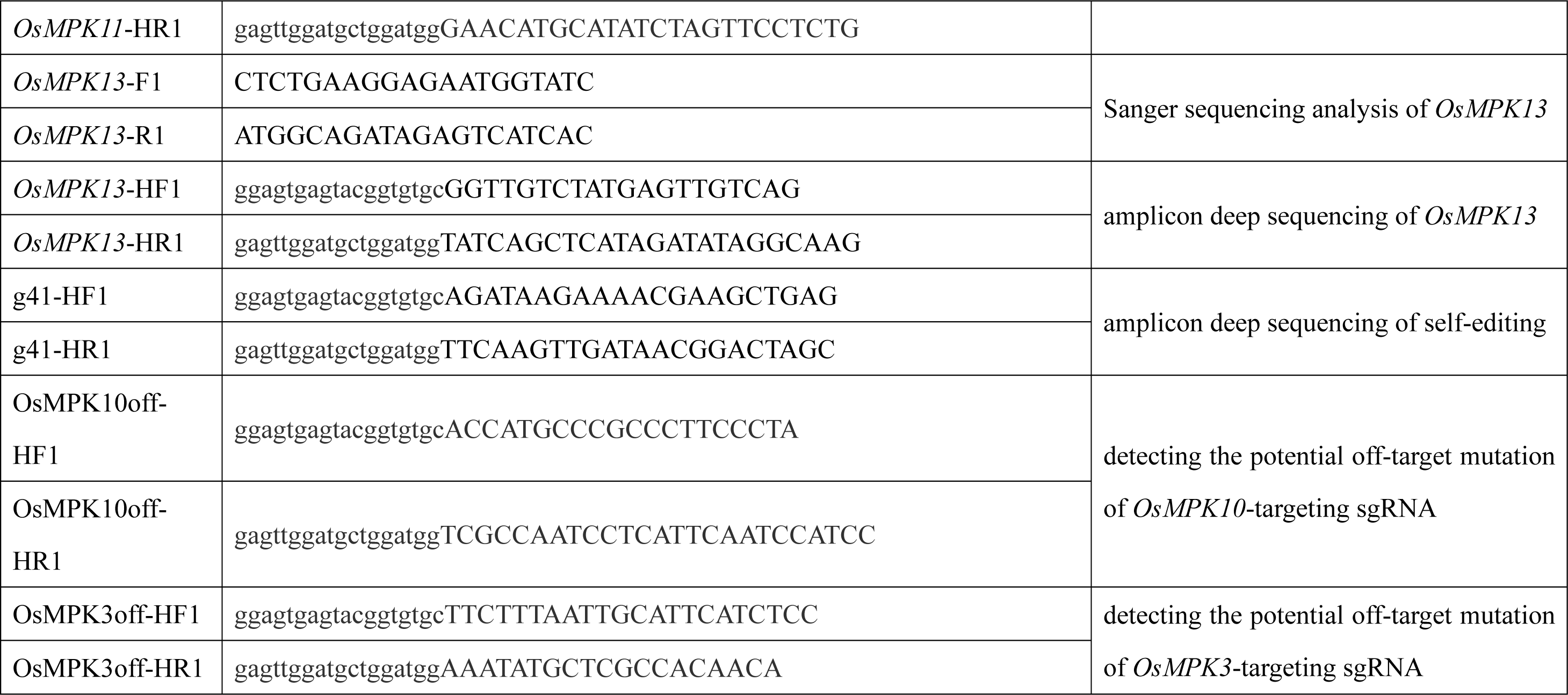
List of oligonucleotides used in this study.

## References

Adikusuma F, Lushington C, Arudkumar J, Godahewa GI, Chey YCJ, Gierus L, Piltz S, Geiger A, Jain Y, Reti D, Wilson LOW, Bauer DC, Thomas PQ. Optimized nickase- and nuclease-based prime editing in human and mouse cells. Nucleic Acids Res. 2021:49(18):10785–10795. 10.1093/nar/gkab792.

Aird EJ, Lovendahl KN, St Martin A, Harris RS, Gordon WR. Increasing Cas9-mediated homology-directed repair efficiency through covalent tethering of DNA repair template. Commun Biol. 2018:1:54. 10.1038/s42003-018-0054-2.

Anzalone AV, Gao XD, Podracky CJ, Nelson AT, Koblan LW, Raguram A, Levy JM, Mercer JAM, Liu DR. Programmable deletion, replacement, integration and inversion of large DNA sequences with twin prime editing. Nat Biotechnol. 2022:40(5):731–740. 10.1038/s41587-021-01133-w.

Anzalone AV, Randolph PB, Davis JR, Sousa AA, Koblan LW, Levy JM, Chen PJ, Wilson C, Newby GA, Raguram A, Liu DR. Search-and-replace genome editing without double-strand breaks or donor DNA. Nature. 2019:576(7785):149-157. 10.1038/s41586-019-1711-4.

Bill Kim Y, Pierce EB, Brown M, Peterson BA, Sanford D, Fear J, Nicholl D, San Pedro E, Reynolds GM, Hunt JE, Schwark DG, Jali S, Graham N, Cesarz Z, Chapman TAL, Watts JM, Hummel AW. A novel mechanistic framework for precise sequence replacement using reverse transcriptase and diverse CRISPR-Cas systems. bioRxiv. 2022. 10.1101/2022.12.13.520319.

Chatterjee P, Jakimo N, Jacobson JM. Minimal PAM specificity of a highly similar SpCas9 ortholog. Sci Adv. 2018:4(10):eaau0766. 10.1126/sciadv.aau0766.

Chen PJ, Liu DR. Prime editing for precise and highly versatile genome manipulation. Nat Rev Genet. 2023:24(3):161–177. 10.1038/s41576-022-00541-1.

Chen PJ, Hussmann JA, Yan J, Knipping F, Ravisankar P, Chen PF, Chen C, Nel son JW, Newby GA, Sahin M, Osborn MJ, Weissman JS, Adamson B, Liu DR. Enhanced prime editing systems by manipulating cellular determinants of editing outcomes. Cell. 2021:184(22):5635–5652.e5629. 10.1016/j.cell.2021.09.018.

Doman JL, Pandey S, Neugebauer ME, An M, Davis JR, Randolph PB, McElroy A, Gao XD, Raguram A, Richter MF, Everette KA, Banskota S, Tian K, Tao YA, Tolar J, Osborn MJ, Liu DR. Phage-assisted evolution and protein engineering yield compact, efficient prime editors. Cell. 2023:186(18):3983-4002.e3926. 10.1016/j.cell.2023.07.039.

Ferreira da Silva J, Tou CJ, King EM, Eller ML, Rufino-Ramos D, Ma L, Cromwell CR, Metovic J, Benning FMC, Chao LH, Eichler FS, Kleinstiver BP. Click editing enables programmable genome writing using DNA polymerases and HUH endonucleases. Nat Biotechnol. 2024. 10.1038/s41587-024-02324-x.

Gao C. Genome engineering for crop improvement and future agriculture. Cell. 2021:184(6):1621–1635. 10.1016/j.cell.2021.01.005.

Gibson TJ, Seiler M, Veitia RA. The transience of transient overexpression. Nat Methods. 2013:10(8):715–721. 10.1038/nmeth.2534.

Hong C, Han JH, Hwang GH, Bae S, Seo PJ. Genome-wide *in-locus* epitope tagging of Arabidopsis proteins using prime editors. BMB Rep. 2024:57(1):66-70. 10.5483/BMBRep.2023-0055.

Hua K, Jiang Y, Tao X, Zhu J. Precision genome engineering in rice using prime editing system. Plant Biotechnol J. 2020:18(11):2167–2169. 10.1111/pbi.13395.

Jiang T, Zhang XO, Weng Z, Xue W. Deletion and replacement of long genomic sequences using prime editing. Nat Biotechnol. 2022a:40(2):227–234. 10.1038/s41587-021-01026-y.

Jiang Y, Chai Y, Lu M, Han X, Lin Q, Zhang Y, Zhang Q, Zhou Y, Wang X, Gao C, Chen Q. Prime editing efficiently generates W542L and S621I double mutations in two *ALS* genes in maize. Genome Biol. 2020:21(1):257. 10.1186/s13059-020-02170-5.

Jiang Y, Chai Y, Qiao D, Wang J, Xin C, Sun W, Cao Z, Zhang Y, Zhou Y, Wang XC, Chen QJ. Optimized prime editing efficiently generates glyphosate-resistant rice plants carrying homozygous TAP-IVS mutation in *EPSPS*. Mol Plant. 2022b:15(11):1646–1649. 10.1016/j.molp.2022.09.006.

Kumar J, Char SN, Weiss T, Liu H, Liu B, Yang B, Zhang F. Efficient protein tagging and cis-regulatory element engineering via precise and directional oligonucleotide-based targeted insertion in plants. Plant Cell. 2023:35(8):2722–2735. 10.1093/plcell/koad139.

Li E, Liu H, Huang L, Zhang X, Dong X, Song W, Zhao H, Lai J. Long-range interactions between proximal and distal regulatory regions in maize. Nat. Commun. 2019:10(1):2633. 10.1038/s41467-019-10603-4.

Li H, Zhu Z, Li S, Li J, Yan L, Zhang C, Ma Y, Xia L. Multiplex precision gene editing by a surrogate prime editor in rice. Mol Plant. 2022a:15(7):1077-1080. 10.1016/j.molp.2022.05.009.

Li J, Chen L, Liang J, Xu R, Jiang Y, Li Y, Ding J, Li M, Qin R, Wei P. Development of a highly efficient prime editor 2 system in plants. Genome Biol. 2022b:23(1):161. 10.1186/s13059-022-02730-x.

Li J, Ding J, Zhu J, Xu R, Gu D, Liu X, Liang J, Qiu C, Wang H, Li M, Qin R, Wei P. Prime editing-mediated precise knockin of protein tag sequences in the rice genome. Plant Commun. 2023a:4(3):100572. 10.1016/j.xplc.2023.100572.

Li X, Zhang G, Huang S, Liu Y, Tang J, Zhong M, Wang X, Sun W, Yao Y, Ji Q, Wang X, Liu J, Zhu S, Huang X. Development of a versatile nuclease prime editor with upgraded precision. Nat Commun. 2023b:14(1):305. 10.1038/s41467-023-35870-0.

Liang R, He Z, Zhao KT, Zhu H, Hu J, Liu G, Gao Q, Liu M, Zhang R, Qiu JL, Gao C. Prime editing using CRISPR-Cas12a and circular RNAs in human cells. Nat Biotechnol. 2024. 10.1038/s41587-023-02095-x.

Lin Q, Jin S, Zong Y, Yu H, Zhu Z, Liu G, Kou L, Wang Y, Qiu J, Li J, Gao C. High-efficiency prime editing with optimized, paired pegRNAs in plants. Nat Biotechnol. 2021:39(8):923–927. 10.1038/s41587-021-00868-w.

Lin Q, Zong Y, Xue C, Wang S, Jin S, Zhu Z, Wang Y, Anzalone AV, Raguram A, Doman JL, Liu DR, Gao C. Prime genome editing in rice and wheat. Nat Biotechnol. 2020:38(5):582–585. 10.1038/s41587-020-0455-x.

Liu B, Dong X, Zheng C, Keener D, Chen Z, Cheng H, Watts JK, Xue W, Sontheimer EJ. Targeted genome editing with a DNA-dependent DNA polymerase and exogenous DNA-containing templates. Nat. Biotechnol. 2023. 10.1038/s41587-023-01947-w.

Liu G, Lin Q, Jin S, Gao C. The CRISPR-Cas toolbox and gene editing technologies. Mol Cell. 2022:82(2):333–347. 10.1016/j.molcel.2021.12.002.

Liu X, Wang Y, Wang H, He Y, Song Y, Li Z, Li M, Wei C, Dong Y, Xue L, Zhang J, Zhu J, Wang M. Generating herbicide resistant and dwarf rice germplasms through precise sequence insertion or replacement. Plant Biotechnol J. 2024:22(2):293–295. 10.1111/pbi.14225.

Lu Y, Ronald PC, Han B, Li J, Zhu J. Rice protein tagging project: A call for international collaborations on genome-wide *in-locus* tagging of rice proteins. Mol Plant. 2020a:13(12):1663–1665. 10.1016/j.molp.2020.11.006.

Lu Y, Tian Y, Shen R, Yao Q, Wang M, Chen M, Dong J, Zhang T, Li F, Lei M, Zhu JK. Targeted, efficient sequence insertion and replacement in rice. Nat Biotechnol. 2020b:38(12):1402–1407. 10.1038/s41587-020-0581-5.

Nambiar TS, Baudrier L, Billon P, Ciccia A. CRISPR-based genome editing through the lens of DNA repair. Mol Cell. 2022:82(2):348–388. 10.1016/j.molcel.2021.12.026.

Nelson JW, Randolph PB, Shen SP, Everette KA, Chen PJ, Anzalone AV, An M, Newby GA, Chen JC, Hsu A, Liu DR. Engineered pegRNAs improve prime editing efficiency. Nat Biotechnol. 2022:40(3):402–410. 10.1038/s41587-021-01039-7.

Ni P, Zhao Y, Zhou X, Liu Z, Huang Z, Ni Z, Sun Q, Zong Y. Efficient and versatile multiplex prime editing in hexaploid wheat. Genome Biol. 2023:24(1):156. 10.1186/s13059-023-02990-1.

Peterka M, Akrap N, Li S, Wimberger S, Hsieh P, Degtev D, Bestas B, Barr J, van de Plassche S, Mendoza Garcia P, Šviković S, Sienski G, Firth M, Maresca M. Harnessing DSB repair to promote efficient homology-dependent and -independent prime editing. Nat Commun. 2022:13(1):1240. 10.1038/s41467-022-28771-1.

Porebski S, Bailey LG, Baum BR. Modification of a CTAB DNA extraction protocol for plants containing high polysaccharide and polyphenol components. Plant Mol Biol Rep. 1997:15(1):8–15. 10.1007/BF02772108.

Ren B, Liu L, Li S, Kuang Y, Wang J, Zhang D, Zhou X, Lin H, Zhou H. Cas9-NG greatly expands the targeting scope of the genome-editing toolkit by recognizing NG and other atypical PAMs in rice. Mol Plant. 2019:12(7):1015-1026. 10.1016/j.molp.2019.03.010.

Schwinn MK, Machleidt T, Zimmerman K, Eggers CT, Dixon AS, Hurst R, Hall MP, Encell LP, Binkowski BF, Wood KV. CRISPR-mediated tagging of endogenous proteins with a luminescent peptide. ACS Chem Biol. 2018:13(2):467–474. 10.1021/acschembio.7b00549.

Shi B, Ni L, Liu Y, Zhang A, Tan M, Jiang M. *OsDMI3*-mediated activation of *OsMPK1* regulates the activities of antioxidant enzymes in abscisic acid signalling in rice. Plant Cell Environ. 2014:37(2):341–352. 10.1111/pce.12154.

Tan J, Zhao Y, Wang B, Hao Y, Wang Y, Li Y, Luo W, Zong W, Li G, Chen S, Ma K, Xie X, Chen L, Liu YG, Zhu Q. Efficient CRISPR/Cas9-based plant genomic fragment deletions by microhomology-mediated end joining. Plant Biotechnol J. 2020:18(11):2161–2163. 10.1111/pbi.13390.

Tang X, Sretenovic S, Ren Q, Jia X, Li M, Fan T, Yin D, Xiang S, Guo Y, Liu L, Zheng X, Qi Y, Zhang Y. Plant prime editors enable precise gene editing in rice cells. Mol Plant. 2020:13(5):667-670. 10.1016/j.molp.2020.03.010.

Tao R, Wang Y, Hu Y, Jiao Y, Zhou L, Jiang L, Li L, He X, Li M, Yu Y, Chen Q, Yao S. WT-PE: Prime editing with nuclease wild-type Cas9 enables versatile large-scale genome editing. Signal Transduct Target Ther. 2022:7(1):108. 10.1038/s41392-022-00936-w.

Tian Y, Zhong D, Shen R, Tan X, Zhu C, Li K, Yao Q, Li X, Zhang X, Cao X, Wa ng P, Zhu JK, Lu Y. Rapid and dynamic detection of endogenous proteins through *in-locus* tagging in rice. Plant Commun. 2024. 10.1016/j.xplc.2024.101040101040.

Walton RT, Christie KA, Whittaker MN, Kleinstiver BP. Unconstrained genome targeting with near-PAMless engineered CRISPR-Cas9 variants. Science. 2020:368(6488):290-296. 10.1126/science.aba8853.

Wang C, Cheng JKW, Zhang Q, Hughes NW, Xia Q, Winslow MM, Cong L. Microbial single-strand annealing proteins enable CRISPR gene-editing tools with improved knock-in efficiencies and reduced off-target effects. Nucleic Acids Res. 2021a:49(6):e36. 10.1093/nar/gkaa1264.

Wang J, He Z, Wang G, Zhang R, Duan J, Gao P, Lei X, Qiu H, Zhang C, Zhang Y, Yin H. Efficient targeted insertion of large DNA fragments without DNA donors. Nat Methods. 2022a:19(3):331–340. 10.1038/s41592-022-01399-1.

Wang L, Kaya HB, Zhang N, Rai R, Willmann MR, Carpenter SCD, Read AC, Martin F, Fei Z, Leach JE, Martin GB, Bogdanove AJ. Spelling changes and fluorescent tagging with prime editing vectors for plants. Front Genome Ed. 2021b:3. 10.3389/fgeed.2021.617553.

Wang M, Yan F, Zhou H. Protocol for targeted modification of the rice genome using base editing. STAR Protoc. 2022b:3(4):101865. 10.1016/j.xpro.2022.101865.

Wang M, Li S, Li H, Song C, Xie W, Zuo S, Zhou X, Zhou C, Ji Z, Zhou H. Genome editing of a dominant resistance gene for broad-spectrum resistance to bacterial diseases in rice without growth penalty. Plant Biotechnol J. 2024:22(3):529–531. 10.1111/pbi.14233.

Wang M, Xu Z, Gosavi G, Ren B, Cao Y, Kuang Y, Zhou C, Spetz C, Yan F, Zhou X, Zhou H. Targeted base editing in rice with CRISPR/ScCas9 system. Plant Biotechnol J. 2020:18(8):1645–1647. 10.1111/pbi.13330.

Wang S, Mao H, Hou L, Hu Z, Wang Y, Qi T, Tao C, Yang Y, Zhang C, Li M, Liu H, Hu S, Chai R, Wang Y. Compact SchCas9 recognizes the simple NNGR PAM. Adv Sci. 2022c:9(4):e2104789. 10.1002/advs.202104789.

Xie X, Ma X, Zhu Q, Zeng D, Li G, Liu YG. CRISPR-GE: A convenient software toolkit for CRISPR-based genome editing. Mol Plant. 2017:10(9):1246–1249. 10.1016/j.molp.2017.06.004.

Xu W, Yang Y, Yang B, Krueger CJ, Xiao Q, Zhao S, Zhang L, Kang G, Wang F, Yi H, Ren W, Li L, He X, Zhang C, Zhang B, Zhao J, Yang J. A design optimized prime editor with expanded scope and capability in plants. Nat Plants. 2022:8(1):45-52. 10.1038/s41477-021-01043-4.

Xu Z, Kuang Y, Ren B, Yan D, Yan F, Spetz C, Sun W, Wang G, Zhou X, Zhou H. SpRY greatly expands the genome editing scope in rice with highly flexible PAM recognition. Genome Biol. 2021:22(1):6. 10.1186/s13059-020-02231-9.

Yan D, Ren B, Liu L, Yan F, Li S, Wang G, Sun W, Zhou X, Zhou H. High-efficiency and multiplex adenine base editing in plants using new TadA variants. Mol Plant. 2021:14(5):722–731. 10.1016/j.molp.2021.02.007.

Yusibov V, Kushnir N, Streatfield SJ. Antibody production in plants and green algae. Annu Rev Plant Biol. 2016:67:669–701. 10.1146/annurev-arplant-043015-111812.

Zheng C, Liu B, Dong X, Gaston N, Sontheimer EJ, Xue W. Template-jumping prime editing enables large insertion and exon rewriting in vivo. Nat Commun. 2023:14(1):3369. 10.1038/s41467-023-39137-6.

Zhong Z, Fan T, He Y, Liu S, Zheng X, Xu Y, Ren J, Yuan H, Xu Z, Zhang Y. An improved plant prime editor for efficient generation of multiple-nucleotide variations and structural variations in rice. Plant Commun. 2024. 10.1016/j.xplc.2024.100976.

Zhou H, Liu B, Weeks DP, Spalding MH, Yang B. Large chromosomal deletions and heritable small genetic changes induced by CRISPR/Cas9 in rice. Nucleic Acids Res. 2014:42(17):10903–10914. 10.1093/nar/gku806.

Zhu H, Li C, Gao C. Applications of CRISPR-Cas in agriculture and plant biotechnology. Nat Rev Mol Cell Biol. 2020:21(11):661–677. 10.1038/s41580-020-00288-9.

Zong Y, Liu Y, Xue C, Li B, Li X, Wang Y, Li J, Liu G, Huang X, Cao X, Gao C. An engineered prime editor with enhanced editing efficiency in plants. Nat Biotechnol. 2022:40(9):1394-1402. 10.1038/s41587-022-01254-w.

